# Ungulate conservation: Lessons from experimental white-lipped peccary management in agricultural-natural landscape mosaics of the Brazilian Cerrado

**DOI:** 10.64898/2026.04.03.716323

**Authors:** Ennio Painkow Neto, Kirsten Mariana Silvius, Gonzalo Barquero, Danilo Carvalho Neves, José Manuel Vieira Fragoso

## Abstract

Animal population control is widely used to mitigate conflicts between wildlife and agriculture worldwide. Structured, monitored removals are rare in South America, however, and their consequences for wildlife populations as well as their effectiveness in reducing crop damage are little understood. Using eight years of data from an experimental white-lipped peccary management program in an agricultural mosaic in the Brazilian Cerrado biome, we assess how structured, non-lethal removals affect both peccary demography and second-crop corn damage. Leslie removal models based on 6,619 captured individuals indicated that cumulative removals to approximately 85% of the initial population strongly reduced peccary abundance, with limited demographic compensation despite fluctuations in reproductive output. Corn crop damage, quantified with satellite imagery, declined over time and was correlated with peccary population size. Interannual variation in population growth and juvenile recruitment was poorly explained by climate, fire, or landscape composition. Source-sink dynamics likely play a role in maintaining healthy populations at the regional scale. Together, these results demonstrate that sustained and monitored ungulate removals can reliably reduce population size and agricultural damage, supporting coexistence between wildlife and food crop production in human-dominated tropical landscapes.

## Introduction

Earth’s current biodiversity crisis unfolds in the Anthropocene, an era in which human population growth, land-use change, and resource demand increasingly shape ecological dynamics at local, regional and planetary scales [1]. Among these pressures, agricultural expansion and intensification stand out as dominant forces restructuring ecosystems, driving habitat loss, species extinctions and cumulative degradation to meet the increasing human demand for food [2,3]. Reconciling biodiversity conservation with food production has become a central challenge for land and resource managers.

Because agriculture now occupies a large fraction of the terrestrial surface, biodiversity conservation increasingly depends not only on protected areas but also on the management of human-dominated landscapes [2,3]. In agricultural frontiers globally, conflicts between wildlife and crop production result from the spatial overlap between mobile animal populations and monocultures interspersed with natural landscapes [4]. The persistence and even abundance of wildlife populations, including endangered species, in these landscapes highlights their potential for conservation, but also the need for public policy and technical solutions to balance biodiversity conservation with human food security. This is especially true in South America, where over the past two decades mechanized agricultural expansion far surpasses the global average [5], leading to increasing human-wildlife conflicts in the mega-biodiverse countries of the region that must be addressed at landscape scales.

Wildlife population management, including population control through culling, is a widely used response to mitigate economic losses and maintain social tolerance toward biodiversity in human-dominated landscapes [4,6]. However, the demographic consequences of such management interventions remain poorly understood for threatened, mobile, and social species [7], including ungulates. Studies in temperate regions show that high mobility, strong social cohesion, and elevated reproductive potential can generate compensatory responses to culling in ungulates that undermine sustained population reduction, particularly in open and highly connected landscapes where removed individuals may be rapidly replaced [8,9].

Much less information is available on compensatory responses in Neotropical ungulate population dynamics [10]. In South America, agricultural expansion currently occurs in forested or woodland areas. Research on human-wildlife conflicts in these agricultural frontier landscapes is scarce. The population biology and dynamics data available to support management for some of these species are valid only in the context of subsistence or small-scale commercial hunting in natural habitats [11,12,13] and rely heavily on the precautionary principle [14].

An additional critical knowledge gap for wildlife management in agricultural frontiers is that while most control strategies implicitly assume a proportional relationship between animal abundance and realized crop damage, explicit evaluations of this assumption are rare globally. Few studies have jointly examined temporal variation in population size and agricultural losses using quantitative data [15], even though damage reduction is often the primary objective of management interventions. This lack of information limits our ability to assess whether population control delivers the outcomes expected by stakeholders, separate from its impact on population dynamics.

Management outcomes are further shaped by environmental context. For large herbivores in temperate systems, interactions among density, climate variability, and habitat condition influence population growth and recruitment, thereby modulating the effectiveness of harvest or culling regimes [16,17,18]. Empirical work on threatened species such as woodland caribou (*Rangifer tarandus caribou*) shows that demographic variables can vary substantially among years in time-lagged response to climatic conditions, even at low densities, although climate alone often explains only a portion of this variability and its effect may be a consequence of habitat marginality [19]. These findings underscore the need to interpret management outcomes within broader environmental contexts. In a review, [20] concluded that for species in the temperate and subtropical zones, and African woodlands and grasslands, primary productivity, itself driven by environmental variation, is a controlling factor in population growth rates.

Empirical studies from African savannas further show that rainfall strongly regulates ungulate population dynamics across species and temporal scales, often overriding density-dependent effects [21]. In contrast, demographic responses of tropical forest ungulates to environmental variability remain poorly documented, with limited information on vital rates, population growth, and carrying capacity (K), which constrains predictions of how environmental change and harvesting interact to influence population persistence [22].

Within this context, the white-lipped peccary (*Tayassu pecari*), a wide-ranging and highly social ungulate of Neotropical forests and savanna-woodlands, is a case in point and may also provide a valuable model system. From a population dynamic perspective, it is of particular interest because the species is known to exhibit strong population cycles in continuous, unbroken forest [10], cycles which could interact with management interventions. The species plays key functional roles through seed predation and soil disturbance [23,24,25] and is nationally and globally listed as threatened [26,27]. Nevertheless, white-lipped peccaries (WLPs) have adapted to exploit large-scale agricultural landscapes. In Brazil they are among the main vertebrate sources of damage to the corn crop [5,28,29,30]. In the region surrounding Emas National Park (ENP), a large federal conservation area in the Brazilian Cerrado, WLP damage has been estimated to affect on average 12.9% of corn fields, resulting in substantial economic losses for producers [30].

Historically affected by habitat loss, hunting, and disease [31,32], WLP populations are thus now increasingly exposed to conflict with industrial agriculture [30,33]. In several regions, recurrent crop losses have led to informal and unregulated lethal control by producers [28,29]. Because these actions are typically implemented without demographic monitoring, they raise concerns that conflict-driven persecution may fail to produce durable reductions in agricultural damage while accelerating population declines of a threatened species.

Despite the scale of these conflicts, long-term quantitative evaluations of structured, science-based removal programs remain rare for any ungulates and are non-existent for WLP. We therefore lack empirical understanding of how sustained removals alter population demography in WLP and other tropical ungulates, how changes in abundance translate into crop damage through time, and how these responses interact with climatic variability and landscape configuration in highly connected agricultural frontiers.

Here, we analyze an eight-year time series from a mosaic of agricultural and natural habitat patches in the Cerrado biome, a threatened biodiversity hotspot [34], to quantify the demographic and agricultural consequences of a government-approved WLP removal program designed to reduce damage to corn crops. We ask 1) whether cumulative removals reduce population size and alter age structure, 2) whether temporal variation in crop damage tracks changes in population size under sustained management, and 3) how interannual variation in population growth and juvenile recruitment relates to precipitation, temperature, fire, and landscape composition. By integrating demographic, agricultural, and environmental data for a threatened social ungulate, our study provides a rare long-term evaluation of wildlife management in an active agricultural frontier and offers general insights into how and when population management can mitigate conflict between wildlife and food production in human-dominated landscapes.

## Material and methods

### Study area

The study was conducted in the central-western region of Brazil, at the junction of the states of Goiás, Mato Grosso and Mato Grosso do Sul, on the southern portion of ENP and surrounding properties (central coordinates: 18.312087° S, 52.892198° W; Figure 1). The original vegetation was dominated by native grasslands, wetlands, and savanna–woodland formations, interspersed with patches of native forest [33,35,36] (Figure 1). The climate is tropical seasonal, with a rainy season from October to April and a dry season from May to September; mean annual precipitation is ∼1,100 mm and mean annual temperature ranges from 22 to 24 °C [37].

**Figure 1.**
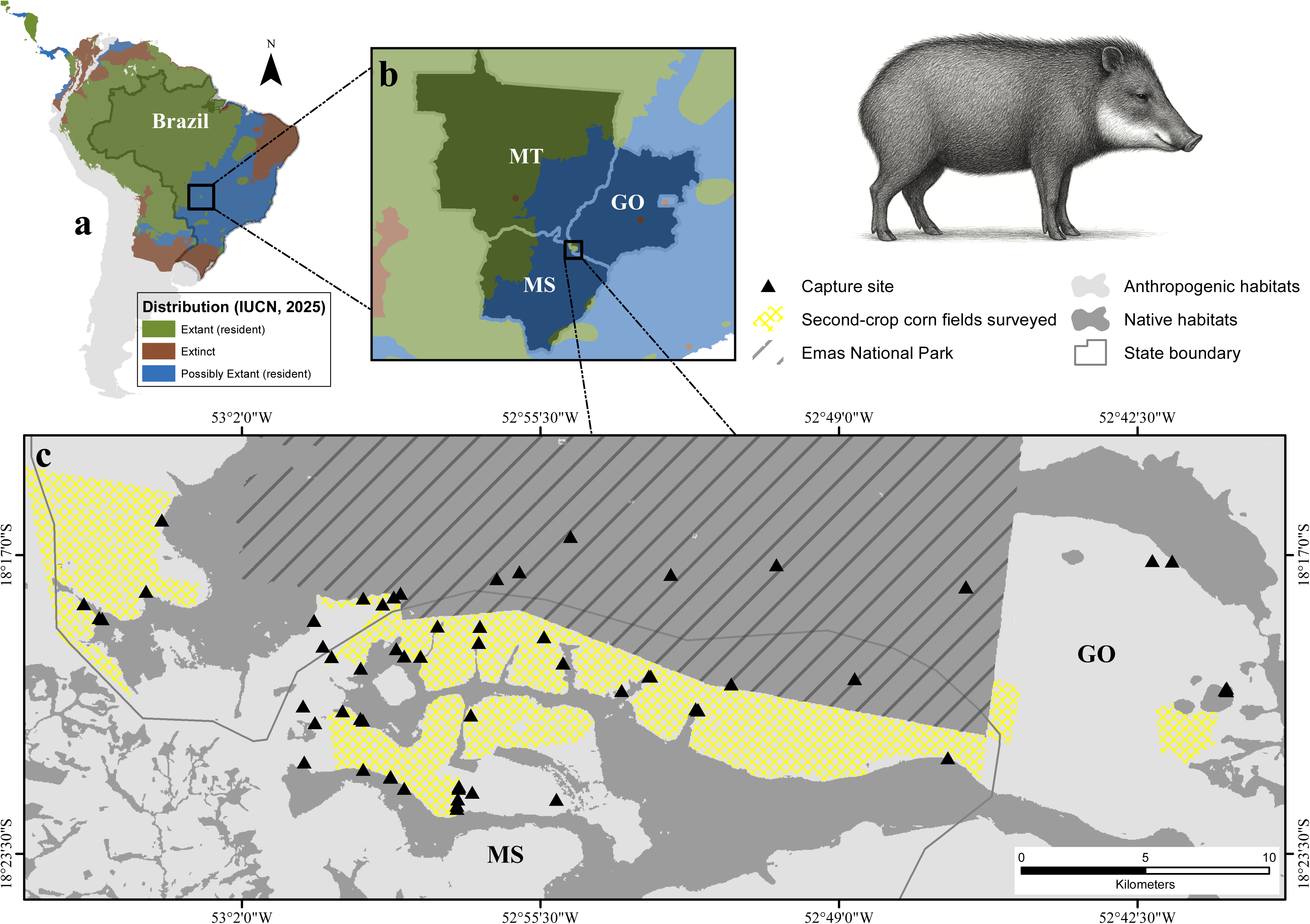
Spatial context of the study system and landscape configuration. (a) Geographic distribution of the WLP in South America according to the International Union for Conservation of Nature (IUCN) [27], with the location of the study area indicated. (b) Regional setting of the study area within the southern Cerrado, central Brazil, encompassing the states of Goiás, Mato Grosso, and Mato Grosso do Sul. (c) Local configuration of the agricultural–natural mosaic, showing the spatial arrangement of native and anthropogenic habitats, capture locations, and second-crop corn fields surveyed for crop damage.

Established in 1961, the 132,000-ha ENP preserves one of the largest remaining blocks of native Cerrado vegetation and protects the headwaters of several river systems [35]. Surrounding landscapes, however, underwent rapid land-use change in subsequent decades. Historically, native vegetation in this region was first converted to extensive cattle pastures, and from the 1970s onwards federal development programs (e.g., PRODECER and POLOCENTRO) accelerated the expansion of mechanized croplands [28]. The current agricultural matrix is largely dominated by large-scale agriculture, particularly soybean, corn, sugarcane and cotton, with pastures now residual elements in the landscape. Over recent decades, agricultural systems have been increasingly intensified and reorganized through the widespread adoption of no-till systems, as well as changes in cropping calendars and soybean cultivar cycles, largely driven by disease management (notably Asian soybean rust), which shifted soybean planting to the beginning of the rainy season and shortened its growing period. This temporal reorganization advanced the soybean harvest from late autumn to mid-summer, thereby enabling the widespread expansion of corn cultivation as a second crop (henceforth “second-crop corn”) within the same agricultural year [38,39,40]. In this region, the agricultural calendar is now characterized by two windows: a main season (October–February) dominated by soybean and a second season (February–July) dominated by second-crop corn.

### Capture and management

Between 2017 and 2024 we captured WLP herds at multiple sites in the southern portion of ENP and in eight surrounding farms (Figure 1) as part of a pilot population control project authorized by SISBIO 87199, implemented under Technical Cooperation Agreement no. 02/2017/CR10/CENAP/ICMBio, and funded by the participating landowners. Herds had been tracked since 2019 and their home ranges mapped (Figure S1); herds known to use only the park interior were not captured for the study presented here [30,33]. Capture sites were selected based on signs of frequent use (e.g., wallows, trails and resting areas). WLPs were trapped in ∼140 m² corral traps baited with corn and mineral salt and fitted with remotely or mechanically triggered gates, which reduced escape and allowed the capture of most herd members [30,33]. Capture events were timed to reduce WLP population size prior to ripening of second-crop corn, when peak damage typically occurs. Operations were conducted preferentially during cooler periods of the day to minimize thermal and behavioral stress.

The project followed an adaptive management design with contrasting strategies implemented across years (Figure 1). In 2017, capture operations were conducted exclusively for marking and individual identification, serving to establish the initial population size for demographic analyses rather than to mitigate crop damage. All animals were released immediately after capture. In 2019 and 2024, captured herds were held in fenced facilities throughout the second-crop corn season (up to 9 months), then released after harvest in the same region. This temporary captivity was implemented to test whether removing herds only during the critical period of corn availability would reduce damage while maintaining population integrity. For population growth rate analysis, these animals are counted as remaining in the population.

In 2018, 2020, 2021, and 2023, all captured individuals were permanently removed and transferred to licensed commercial breeding facilities in Goiás state (SEMAD permits 719332). These facilities operate under an ex-situ conservation and sustainable use framework, in which wild-caught founders are maintained as breeding stock and only captive-born offspring are eligible for regulated harvest.

### Age classification

During annual captures from 2017 to 2024, we recorded the age class (juvenile or adult) for each individual, together with the total number of WLPs captured per year. Age classification followed a standardized protocol throughout the entire period to ensure temporal comparability. “Adults” were defined as fully black, clearly reproductive animals and “juveniles” as lighter-colored individuals that were still associating closely with their mothers, often suckling and not yet reproductive. For analysis, records were aggregated by year to obtain annual counts of juveniles and total individuals. Individuals for which age class could not be assigned unambiguously were excluded from the calculation of annual proportions. Sex was not documented during these annual captures, only during the three years when animals were marked and returned to the population and is therefore not included in the demographic analysis.

### Population size estimation by removal (Leslie model)

We defined a capture session (≥ 3 per year) as the continuous period during which corral traps were installed and monitored; their duration and the number of active traps varied among years, reflecting behavioral responses of herds to trapping [41]. We estimated annual population size of WLPs between 2017 and 2024 using the classic removal-based Leslie regression model [42]. For each year, we calculated, at the capture-session level: (i) the number of individuals captured, (ii) capture effort (trap-active days × number of traps), (iii) catch per unit effort (CPUE, individuals per trap day) and (iv) cumulative catch. CPUE for a given session was calculated as the total number of WLPs captured divided by the number of trap-days [43].

Following [44], we regressed session-level *CPUE* 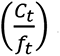 against cumulative catch up to each session (*K_t_*) within each year. Under the standard assumptions of removal models – demographic and geographic closure during sampling, constant catchability and equal capture probability among individuals [45] – this relationship is expected to be linear and negative as the population is progressively depleted:

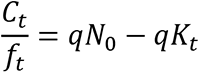

where *C_t_* is the number of individuals captured in session *t*, *f_t_* is the corresponding effort, *K_t_* is cumulative catch up to session *t, q* is the catchability coefficient and *N*_0_is the initial population size. The intercept of the regression (*qN*_0_) and its slope (−*q*) were estimated by ordinary least squares, and annual population size was obtained as 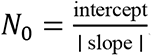. Confidence intervals for *N*_0_ were derived from the variance of the regression parameters following [46].

### Assessment of crop damage in second-crop corn

To quantify crop damage attributed to WLPs between 2017 and 2024, we analyzed satellite imagery for eight farms where management actions were implemented (approximately 10,800 hectares; Figure 1). Damage assessments were conducted at the level of agricultural fields (hereafter “fields”), which represent the primary spatial units of crop production.

Within each field, areas affected by WLPs were identified based on confirmed presence and characteristic damage patterns and were delineated as minimum convex polygons (MCPs) encompassing contiguous patches of impacted vegetation (see Figure S2). These polygons represent the spatial extent of crop loss within each field and were used as the basis for subsequent quantification of damage.

For each year, damage analyses were restricted to fields where WLP damage was confirmed during the second-crop corn season. Fields without evidence of damage in that year were not included in the analysis. Therefore, the area for which damage was quantified varies from year to year. Importantly, the mapped units do not represent the full extent of the farms or all planted fields, but only the fields where WLPs were present during the respective growing season.

Satellite imagery was selected to coincide with the reproductive stage of second-crop corn, typically between late April and late May, when plants exhibit high leaf area and greenness. This phenological window maximizes spectral contrast between intact corn and damaged or exposed soil. Due to variable cloud cover, image acquisition dates could not be fully standardized across years. We used Sentinel-2A/B imagery (10 m spatial resolution) for 2017–2019 and CBERS-4A imagery (8 m multispectral and 2 m panchromatic) for 2020–2024, which were pan-sharpened to enhance spatial resolution [47,48].

Within each selected scene, fields with confirmed WLP damage were identified, and damaged areas within each field were manually delineated as MCPs, which defined the inspected area used to quantify crop loss (Figure S2). The Normalized Difference Vegetation Index (NDVI) was then calculated within these polygons to characterize vegetation condition [48,49]. NDVI rasters were classified using scene-specific thresholds, adjusted individually to maximize discrimination between actively growing corn and exposed soil. Although NDVI values around 0.3 are commonly reported as indicative of bare soil or severely degraded vegetation [48,50], no fixed threshold was imposed across images. Instead, thresholds were tuned based on the NDVI distribution of each scene, reflecting variation in illumination, sensor characteristics, and atmospheric conditions.

Under these conditions, undamaged corn consistently exhibited high or saturated NDVI values, often exceeding 0.8, whereas damaged patches were characterized by very low NDVI values associated with exposed soil and destroyed plants. Pixels classified as damaged were converted to a binary layer, and damaged area was calculated by multiplying the number of damaged pixels by pixel area (25 m² for Sentinel-2 and 4 m² for CBERS-4A). For each field, crop damage was expressed as the proportion of the inspected area classified as damaged, where the inspected area corresponds to the MCP delineating the extent of WLP damage within the field (Figure S2; and see Table S1 for details on area inspected in each farm).

Attribution of damage to WLPs was supported by multiple independent lines of evidence. Farm owners, workers, and agronomists in the region routinely monitor crop fields on a weekly or daily basis due to the economic importance of second-crop corn. WLP damage is well known locally and has been documented in previous studies and technical reports [28,30,33,51].

Damage patterns attributed to WLPs are distinctive, as herds typically move along machinery traffic lanes created by self-propelled sprayers and consume corn plants along these corridors, producing large, irregular patches of destruction that are readily detectable in satellite imagery. Recent studies using UAV-based RGB and multispectral imagery further demonstrate that damage caused by large mammals such as Suiformes can be reliably distinguished from other sources of crop loss based on spatial patterns and vegetation signatures [52].

Other large herbivores or omnivores occur at low abundances in the study region [53] and thus do not produce damage at spatial scales comparable to those caused by WLPs. Alternative sources of crop loss were readily distinguishable in satellite imagery. Planting failures typically produced regular geometric patterns aligned with sowing direction, whereas areas affected by excess soil moisture formed persistent scars that were visible across multiple years, including during periods without cultivation. Such features were excluded from damage classification.

Fields where second-crop corn was not planted, where image interpretation was not possible due to cloud cover, or where access by WLPs was effectively prevented were not considered in the definition of the analytical dataset, as damage was quantified exclusively for fields with confirmed WLP activity.

### Environmental covariates

Precipitation and temperature were obtained from the same fixed weather station at Mineiros, Goiás each year, but fire and land-cover variables were extracted only for the area actually used by the captured herds (Figure S3). Therefore both the total area assessed and the landscape characteristics varied from year to year

#### Climate

Annual precipitation (mm) and mean annual temperature (°C) were derived from the time series of the INMET/BDMEP Mineiros weather station located in a municipality within the study region, which provided the most continuous and stable records for the period analyzed. For each calendar year (January–December), we calculated total annual precipitation and mean annual temperature [37].

#### Fire

Annual burned area data was obtained from the federal government’s MapBiomas Fire Collection 4 database. This database provides annual and monthly maps of burned scars for all of Brazil (1985–2024) based on Landsat mosaics at 30 m resolution. The data is documented in official ATBDs and methodological notes [36]. For each year from 2017 to 2024, we aggregated monthly burn masks into a single annual layer (presence of fire in any month of the year) and calculated the fraction of burned area within the area of interest. This was defined annually as the sum of circular buffers with a radius of 5.24 km centered on each capture site for that year (with buffer overlap areas excluded from the sum; see Fig S3), a value that approximates the radius of a circle with an area equivalent to the 95% fixed kernel home range (∼8,660 ha) of WLP herds in ENP [30,51]. This delineation mirrored the spatial scale of herd space use and tracked the interannual variation in capture locations.

#### Landscape

Land-use and land-cover composition was obtained from MapBiomas Land Cover and Land Use, Collection 10 (30 m resolution, annual series 1985–2024; [36]). For each year from 2017 to 2024, we quantified, within the same area as for fire scars, the percentages of anthropogenic and native cover by aggregating MapBiomas classes. We defined “anthropogenic” as the sum of the following classes: Annual crops (for example, Soybean, Corn, Cotton, Other Temporary Crops), Sugar cane, Pasture, Forest plantation, Urban area, Mosaic of uses and Other non-vegetated areas. We defined “native” as the sum of Forest formation, Savanna formation, Grassland formation, Wetland and River, lake and ocean (MapBiomas national categories).

### Data analysis

#### Effects of management on population size and age structure

To evaluate whether permanent removals affected WLP abundance and age structure, we compiled for each year: (i) the total number of individuals captured, (ii) the number permanently removed from the wild (Rₜ), (iii) the proportion of juveniles among captured individuals and (iv) annual population size (Nₜ) estimated. Years 2017, 2019 and 2024 were “capture–release” years (Rₜ = 0); in 2018 and 2020–2023 all captured individuals were permanently removed (Rₜ > 0). Management intensity was expressed as cumulative removals up to year t (cumulative removalₜ = ΣRₖ, k ≤ t).

We then fitted two simple linear models with Gaussian error: (1) Nₜ ∼ cumulative removalₜ and (2) juvenile proportionₜ ∼ cumulative removalₜ. For both models we inspected residuals for normality and homoscedasticity and report regression coefficients (estimate ± SE), F statistic, degrees of freedom, R² and P-values.

#### Linking population size to crop damage

To evaluate whether population size was associated with reduced agricultural damage, we compiled an annual time series (2017–2024) combining (i) population size estimated (Nₜ) and (ii) crop damage quantified as the proportion of inspected corn area (see methods) destroyed by WLPs in each of the eight farms that same year.

For statistical inference, we related crop damage to population size using linear mixed-effects models with Gaussian error distribution, including farm identity as a random intercept to account for repeated observations and spatial heterogeneity among farms. Model estimates are reported as fixed-effect coefficients ± standard errors, along with F statistics, degrees of freedom, coefficients of determination (marginal and conditional R²), and P-values. Model assumptions were assessed through inspection of residual distributions.

#### Environmental drivers of demographic variation

To assess whether environmental conditions modulated population dynamics in addition to management, we related annual demographic descriptors to climate, fire and land-cover variables extracted from the 5.24 km buffers drawn around each year’s capture sites. We considered two responses: (i) the annual population growth rate 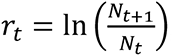, where Nₜ and N₍ₜ₊₁₎ are population size estimated in consecutive years, and (ii) the proportion of juveniles in year t.

For rₜ, we treated year t as the period during which conditions determine the transition from Nₜ to N₍ₜ₊₁₎ and modelled rₜ as a function of environmental variables measured in year t. For juvenile proportionₜ, we assumed that recruitment is shaped mainly by conditions in the preceding year, and related juvenile proportionₜ to environmental variables measured in year t−1.

Given the short time series and strong collinearity among predictors, we used deliberately simple univariate linear models rather than multivariate formulations. For each predictor we fitted two Gaussian models: one with rₜ as the response and another with juvenile proportionₜ.

Predictors for the rₜ models were annual precipitation, mean annual temperature, percentage of burned area, anthropogenic cover and native cover within the focal landscape in year t; the same variables measured in year t−1 were used for the juvenile models. For each model we examined residuals and extracted regression coefficients (estimate ± SE), F, df, R² and P-values, treating these analyses as exploratory tests of whether any single environmental factor showed a consistent association with interannual variation in population growth or juvenile recruitment.

### Statistical software

All statistical analyses were conducted in R version 4.5.2 [54], using *dplyr*, *janitor*, *nlme*, *broom*, *performance*, *ggplot2*, *patchwork*, and *writexl*. Spatial analyses and quantification of agricultural damage were conducted in ArcMap version 10.1 (ESRI, Redlands, CA, USA).

## Results

### General description

#### Effort and captures

Between 2017 and 2024, we captured 6,619 WLPs across the study area (Table 1; Table S*2*). Of these, 1,042 were released immediately, 4,112 were permanently removed, and 1,465 were temporarily removed and released after the second-crop corn harvest. Annual capture effort varied substantially, with an average capture window of 135 ± 31 days and a mean of 827 ± 318 individuals handled per year. The number of capture events ranged from 13 to 54 per year, conducted at 65 capture sites distributed across the southern portion of ENP and surrounding agricultural landscapes (Figure 1). Distances between corral trap locations ranged from 200 to 4,000 m.

**Table 1.** Annual capture effort, management regime and age composition of WLPs in the study area between 2017 and 2024. For each year, we report the start and end dates of the capture window, the total number of individuals captured, the number of juveniles, the number of capture events, and the management type applied. Management type indicates whether captured individuals were released immediately after capture (Capture-release), temporarily held in captivity until corn harvest (Temporary removal), or definitively removed from the wild (Permanent removal). The last two rows report the mean and standard deviation (SD) across years for numeric variables.

| Year | Start Date | End Date | Total individuals captured | Juveniles captured | Number of capture events | Management type |
| --- | --- | --- | --- | --- | --- | --- |
| 2017 | 07/11/2017 | 12/13/2017 | 1,042 | 184 | 26 | Capture-release |
| 2018 | 04/21/2018 | 09/05/2018 | 770 | 54 | 16 | Permanent removal |
| 2019 | 11/28/2018 | 06/11/2019 | 1,122 | 129 | 34 | Temporary removal |
| 2020 | 12/30/2019 | 05/15/2020 | 1,265 | 224 | 54 | Permanent removal |
| 2021 | 02/06/2021 | 06/04/2021 | 544 | 50 | 26 | Permanent removal |
| 2022 | 01/25/2022 | 06/14/2022 | 937 | 216 | 33 | Permanent removal |
| 2023 | 02/08/2023 | 05/18/2023 | 596 | 75 | 18 | Permanent removal |
| 2024 | 04/03/2024 | 07/12/2024 | 343 | 73 | 13 | Temporary removal |
| Total | - | - | 6,619 | 1,005 | 220 | - |
| Mean | - | - | 827 | 126 | 28 | - |
| SD | - | - | 318 | 73 | 13 | - |

#### Proportion of juveniles

Juvenile representation varied markedly among years, ranging from 7.0% to 23.1% of captured individuals (Table 1; Table 2). Over the eight-year study period, juveniles accounted for 15.2% of all WLPs handled (1,005 of 6,619 individuals; Table 1). The mean annual number of juveniles was 126 ± 73, corresponding to an average annual proportion of 15.0 ± 5.8%.

**Table 2.**
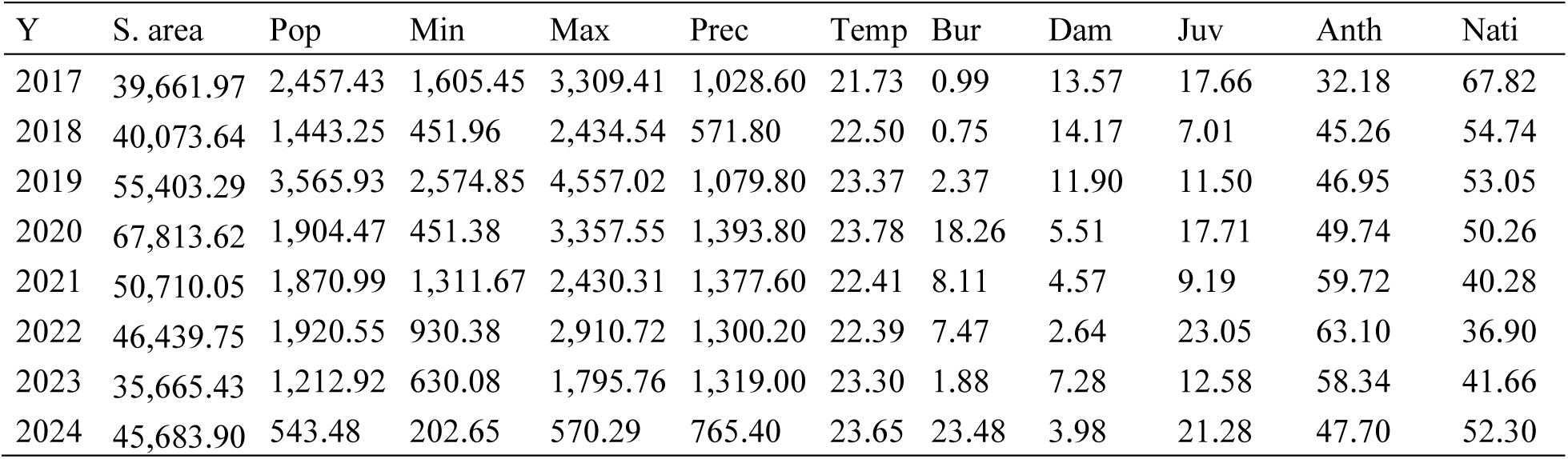
Annual demographic, environmental, landscape, and damage metrics for the WLP population in the study area (2017–2024). Columns report: year (Y); annual study area (S. area, ha); population size estimated by Leslie models (Pop, individuals); lower and upper 95% confidence limits of population size (Min, Max, individuals); annual precipitation (Prec, mm); mean annual temperature (Temp, °C); proportion of the study area affected by fire (Bur, %); mean annual crop damage in second-crop corn (Dam, % of inspected area [MCP] classified as damaged); proportion of juveniles among captured individuals (Juv, %); proportion of anthropogenic land cover (Anth, %); and proportion of native vegetation cover (Nati, %) within the annual study area.

| Y | S. area | Pop | Min | Max | Prec | Temp | Bur | Dam | Juv | Anth | Nati |
| --- | --- | --- | --- | --- | --- | --- | --- | --- | --- | --- | --- |
| 2017 | 39,661.97 | 2,457.43 | 1,605.45 | 3,309.41 | 1,028.60 | 21.73 | 0.99 | 13.57 | 17.66 | 32.18 | 67.82 |
| 2018 | 40,073.64 | 1,443.25 | 451.96 | 2,434.54 | 571.80 | 22.50 | 0.75 | 14.17 | 7.01 | 45.26 | 54.74 |
| 2019 | 55,403.29 | 3,565.93 | 2,574.85 | 4,557.02 | 1,079.80 | 23.37 | 2.37 | 11.90 | 11.50 | 46.95 | 53.05 |
| 2020 | 67,813.62 | 1,904.47 | 451.38 | 3,357.55 | 1,393.80 | 23.78 | 18.26 | 5.51 | 17.71 | 49.74 | 50.26 |
| 2021 | 50,710.05 | 1,870.99 | 1,311.67 | 2,430.31 | 1,377.60 | 22.41 | 8.11 | 4.57 | 9.19 | 59.72 | 40.28 |
| 2022 | 46,439.75 | 1,920.55 | 930.38 | 2,910.72 | 1,300.20 | 22.39 | 7.47 | 2.64 | 23.05 | 63.10 | 36.90 |
| 2023 | 35,665.43 | 1,212.92 | 630.08 | 1,795.76 | 1,319.00 | 23.30 | 1.88 | 7.28 | 12.58 | 58.34 | 41.66 |
| 2024 | 45,683.90 | 543.48 | 202.65 | 570.29 | 765.40 | 23.65 | 23.48 | 3.98 | 21.28 | 47.70 | 52.30 |

#### Population size estimates

Annual population size estimates derived from the Leslie regression model varied from fewer than 600 individuals in the final year of the study to more than 3,500 individuals at its peak, with a mean of 1,865 ± 894 individuals and wide confidence intervals reflecting interannual variability in demographic rates. Detailed annual estimates and associated uncertainty are provided in Table 2.

#### Damage in the second crop corn

Mean crop damage across the eight-year period averaged 8.0 ± 4.6% of the inspected area (Table 2). Damage levels varied markedly among years, ranging from less than 4% in the final years of the study to peaks exceeding 14% in earlier years. Annual estimates represent averages across farms; farm-level variation is reported in the Supporting Information (Table S1).

#### Annual study area and environmental covariates

The annual study area used to quantify burned area and landscape composition varied substantially among years, ranging from 35,665 to 67,814 ha (Table 2). Annual precipitation showed pronounced interannual variability, ranging from less than 600 mm to nearly 1,400 mm, whereas mean annual temperature varied more modestly, spanning approximately 22 to 24 °C (Table 2).

The proportion of burned area within the annual study area also varied widely among years, from values below 1% to peaks exceeding 20%, with an overall mean of 7.9 ± 8.6% (Table 2). Landscape composition exhibited a clear shift through time due to the shifting area in which WLP were captured (increasingly outside of the park in later years; Figure S3), with anthropogenic cover averaging 50.8 ± 10.7% of the study area and increasing across years, while native vegetation showed a complementary decline (Table 2; Figure S4).

### Management and environmental drivers of population dynamics and crop damage

#### Population decline without age-structure shifts

Population size declined with increasing cumulative definitive removals, with a negative relationship indicating a reduction in abundance with increased management intensity (β = −0.38 ± 0.17, R² = 0.46, p = 0.06; Figure 2). The only year with a marked population increase was 2019, one year after the first permanent removals (Figure 2). In contrast, the proportion of juveniles varied among years, with alternating years of low and high reproduction (Figure S4), but showed no consistent change along the gradient of cumulative removals, with effect sizes close to zero. This pattern indicates that population decline was not accompanied by detectable shifts in age structure under sustained management (Figure 2; Table S3).

**Figure 2.**
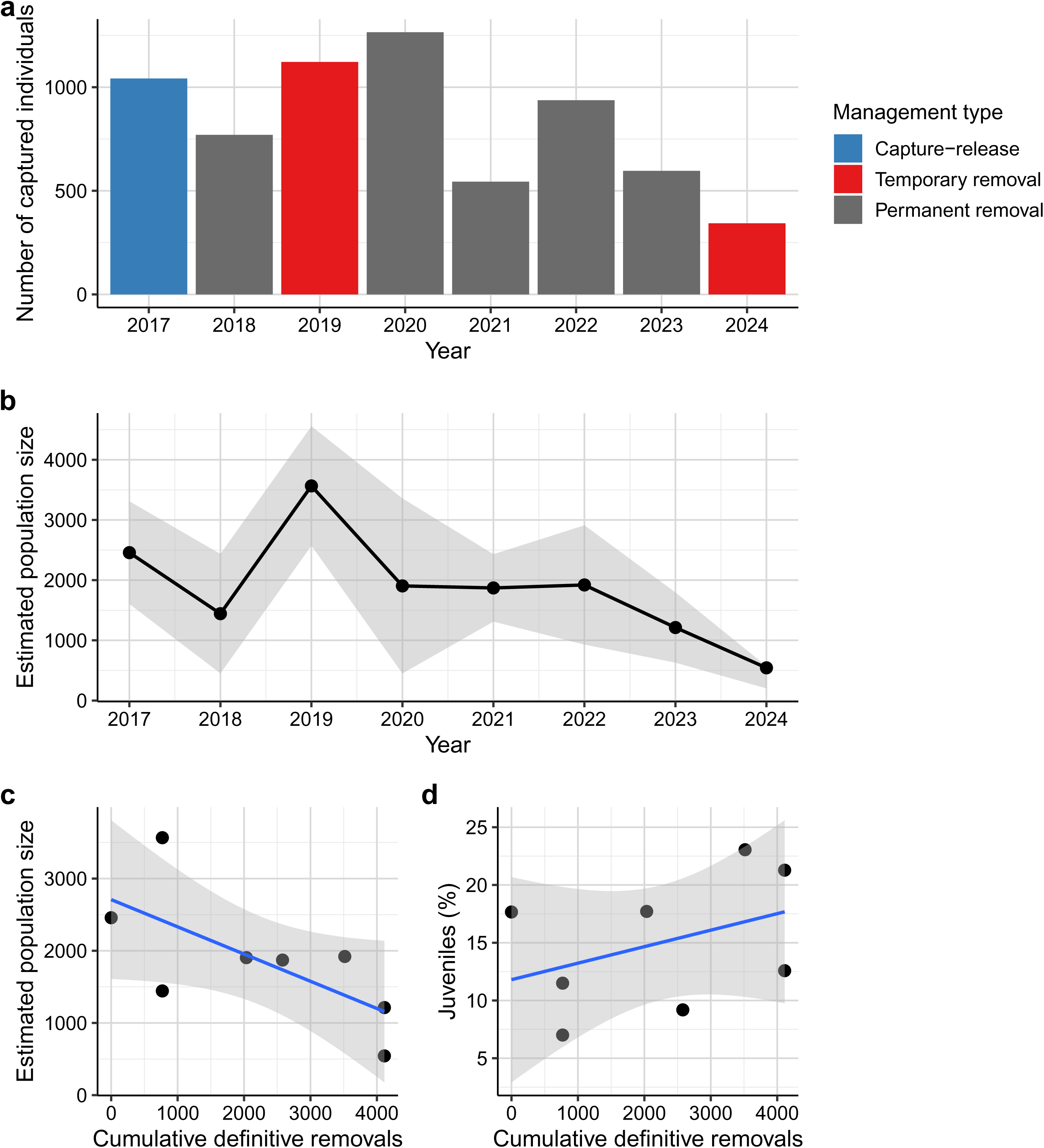
Population responses of WLP to management via definitive removals. (a) Annual number of individuals captured between 2017 and 2024, distinguishing capture–release campaigns (blue bars; 2017), temporary removals from the wild population (red bars; 2019 and 2024), and permanent removals from the wild population (grey bars; 2018, 2020, 2021, 2022, and 2023). (b) Annual population size (N) estimated from the Leslie regression model, with points and connecting line showing point estimates and the shaded band indicating the minimum–maximum range (Leslie “Min” and “Max”). (c) Relationship between estimated population size and the cumulative number of individuals definitively removed over the study period; the blue line shows the fitted simple linear model (N ∼ cumulative removals) and the shaded band its 95% confidence interval. (d) Relationship between the proportion of captured juveniles (%) and cumulative definitive removals; the blue line shows the fitted simple linear model (juveniles ∼ cumulative removals).

#### Crop damage and population size

Crop damage in second-crop corn declined consistently over time, showing a negative temporal trend across the study period (β = −0.010 ± 0.004; Figure 3). After accounting for spatial heterogeneity among farms, crop damage was positively associated with WLP population size, with higher damage observed in years with larger estimated populations (β = 0.022 ± 0.009; Figure 3; Table S3).

**Figure 3.**
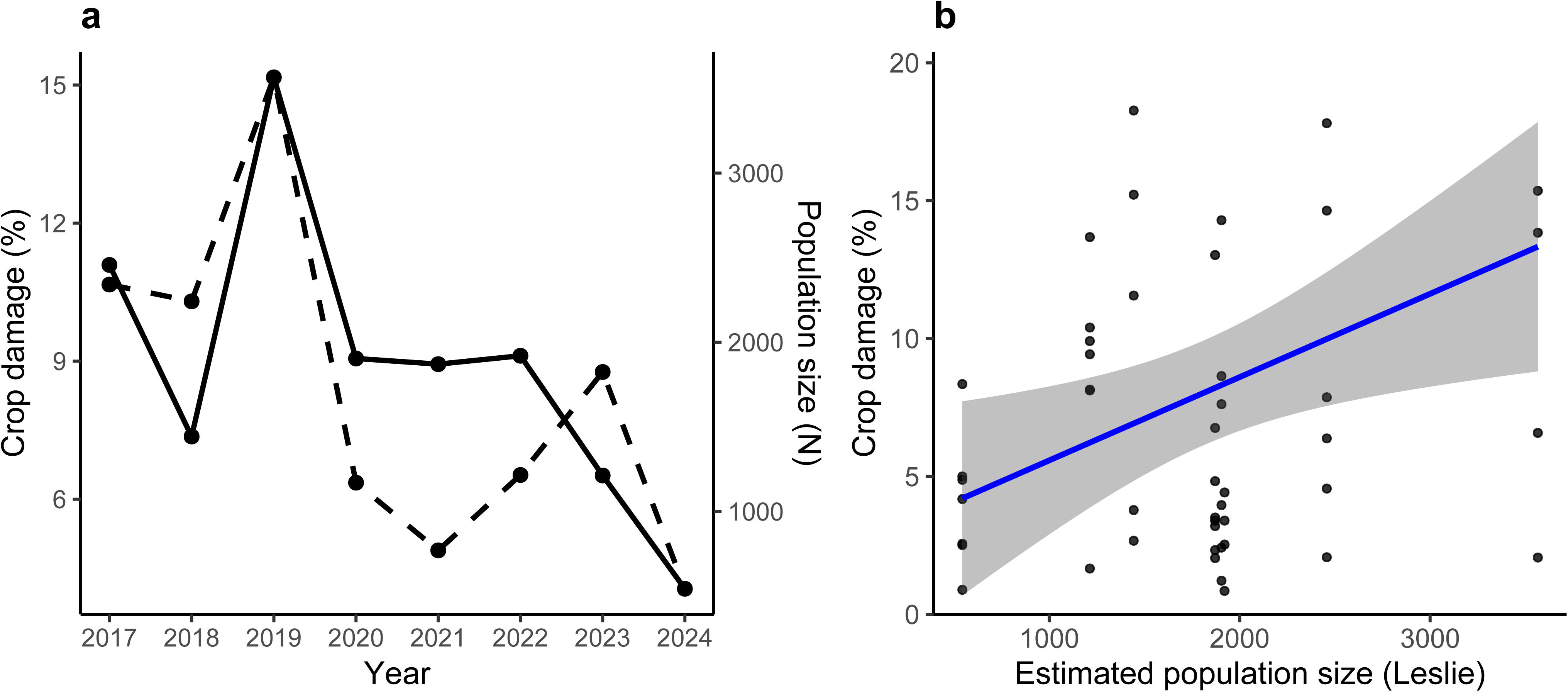
Temporal dynamics and population-damage relationship of WLP crop impacts under management. (a) Annual trajectories of mean crop damage in corn fields (dashed line, left axis) and estimated population size (solid line, right axis) from 2017 to 2024. Population size is shown on a secondary axis to allow direct visual comparison of temporal trends between variables expressed in different units. Both metrics peaked in 2019 and declined thereafter, indicating a concurrent reduction in population size and agricultural damage over time. (b) Relationship between estimated population size derived from the Leslie removal model and crop damage (% of inspected corn area), shown for individual farm–year observations. The solid line represents the fitted linear model, with the shaded area indicating the 95% confidence interval. For clarity of visualization, the y-axis is truncated at 20% crop damage. Crop damage increased with population size, supporting a population-mediated link between management intensity and agricultural losses.

#### Limited environmental effects on population growth and juvenile recruitment

Annual population growth rate showed no clear association with the environmental variables examined. A negative relationship with annual precipitation was observed; however, this pattern was not statistically significant and appeared to be influenced by a single low-rainfall year. Interannual variation in population growth also coincided with changes in management regimes, including years with permanent removals and temporary removals, making it difficult to disentangle potential effects of environmental conditions from those of management actions. Relationships with temperature, burned area, and landscape composition were negligible, with effect sizes near zero (Figure 4; Table S3).

**Figure 4.**
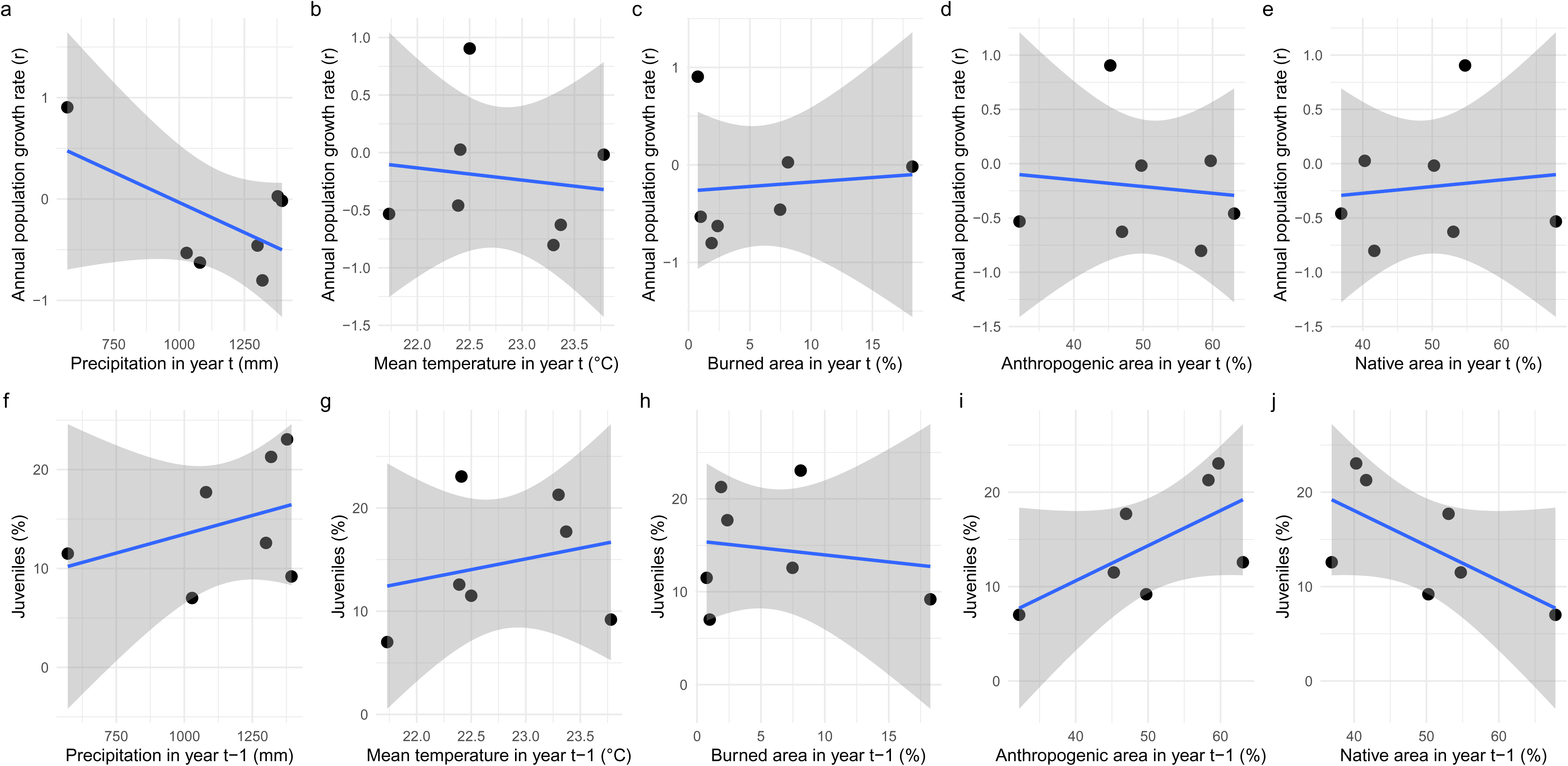
Relationships between annual population growth rate and juvenile proportion of WLP and environmental covariates across the management time series. Panels (a–e): relationships between annual population growth rate 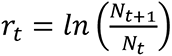, estimated from the Leslie model, and (a) annual precipitation, (b) mean annual temperature, (c) percentage of burned area, (d) percentage of anthropogenic cover and (e) percentage of native cover in year *t*. Panels (f–j): relationships between the proportion of juveniles in year *t* and the same environmental variables measured in the preceding year (*t* − 1), representing the climatic, fire and land-use context under which the juvenile cohort was produced. Black points show annual observations, blue lines, the fitted simple linear regressions and shaded bands the 95% confidence intervals.

Similarly, the proportion of juveniles showed no consistent association with environmental conditions in the preceding year. Relationships with precipitation, temperature and burned area were not statistically significant and showed no clear directional pattern.

Anthropogenic and native land cover exhibited opposite relationships of similar magnitude with proportion of juveniles, but these relationships were also not statistically significant. Overall, environmental variables explained only a limited fraction of interannual variation in both population growth and juvenile recruitment (Figure 4; Table S3).

## Discussion

Our analyses reveal four main findings. First, management through definitive removals led to a marked decline in WLP abundance over eight years, indicating that removal-based control was demographically effective (Figure 2). Second, this decline in abundance was not accompanied by a consistent restructuring of age ratios: the proportion of juveniles fluctuated among years but showed no systematic response to cumulative removals, indicating that removal levels were sufficient to prevent population overshoot from any compensatory recruitment (Figure 2). Third, crop damage in corn fields declined over time under sustained management and covaried with annual population size, indicating that lower WLP abundances were associated with reduced levels of damage (Figure 3). Fourth, interannual variation in population growth and juvenile recruitment was poorly explained by climatic conditions, fire, or landscape composition, suggesting that additional factors (e.g., corn as a food supplement), including management interventions, may contribute to this variability (Figure 2; Figure 4; Figure S4).

Taken together, our data suggest that removal levels were sufficient to prevent population overshoot from compensatory recruitment, but not high enough to raise concern about local population extinction. Continued healthy reproductive levels plus source-sink dynamics among patches in the mosaic demonstrate the conservation value of these mixed landscapes. Animals held in reserve in paddocks have therefore not been needed to restock the population and can be given other destinations.

### Population control and demographic responses

Annual population size estimates ranged from 3,566 individuals in 2019 to 543 in 2024 in the study area (Figure 1), revealing a pronounced decline over the management period (Figure 2). This magnitude of reduction is consistent with findings from systems in which well-structured control programs can effectively depress herbivore densities at broad spatial scales [55], but contrasts with patterns reported for wild boar (*Sus scrofa*), where populations may stabilize or rebound despite substantial removal effort due to compensatory recruitment or recolonization [8,9]. In our system, cumulative definitive removals equivalent to approximately 85% of the estimated population size were associated with a sustained reduction in abundance, indicating that structured removals can be an effective tool to lower local densities of WLP in mixed agricultural-natural landscapes. Given that WLP are a major source of crop damage in these systems [29,30,33,51,56], such reductions in population size have direct implications for mitigating agricultural losses. Such high levels of removal are rarely documented for native ungulate populations, particularly threatened species, under structured and monitored management programs.

WLP life-history traits suggest it has the capacity for compensatory demographic response. In Amazonian forests, where it has been most extensively studied, the species is characterized by moderate reproductive output, delayed maturity, and extended reproductive lifespan, which constrain —but do not eliminate— the potential for rapid density-dependent compensation [57,58,59,60]. In these systems, populations can sustain substantial offtake under certain conditions [11,12,13] and have been documented to exhibit “rapid increases” following periods of low abundance, resulting in pronounced population fluctuations [10].

However, we did not detect evidence of such compensatory responses in our eight-year system. The proportion of juveniles fluctuated among years and, although a slight increasing trend was apparent (Figure S4), this pattern was not statistically supported in relation to cumulative removals, suggesting that management acted largely as additive losses rather than triggering accelerated recruitment at reduced densities.

Offtake in our study was much higher than proposed for sustainable hunting in a natural forested system. This may explain why the sustained high offtake was able to effectively reduce populations without compensatory effects. It also suggests that K in these agricultural–natural mosaics may be much higher than in lower productivity Amazonian forests, likely due to the high availability of energy-rich crops and more predictable resource supply. Population trends in this experimental management initiative must continue to be closely monitored to ensure that the regional population is not threatened.

In this study, removed animals came primarily from herds using forest fragments outside of ENP [30,33], but animals were also removed from the southern region of the park when their home ranges overlapped agricultural areas (Figure S1; Figure S3). When the animals use the corn fields plus native habitat fragments, their dynamics and K may be different, as suggested by the much smaller home range size of herds living outside natural areas, relative to those in ENP [33].

Importantly, the relationship between population size and cumulative removals showed a strong negative effect (slope = −0.377 ± 0.167; R² = 0.46), indicating a substantial reduction in abundance with increasing management intensity. However, statistical support was weak (p = 0.06) probably due to the short time series (n = 8 years), uncertainty in abundance estimation for highly mobile, herd-forming species, and the absence of information on immigration and emigration. Rather than indicating eradication or demographic collapse, the observed pattern is more consistent with population suppression to a lower-density state whose persistence is likely mediated by regional connectivity and recolonization dynamics [33].

From a management perspective, this distinction is critical. The objective of control in agricultural frontiers should not be population elimination, but the maintenance of densities compatible with coexistence. WLPs are a key functional species in Neotropical ecosystems and support biodiversity and livelihoods across their range [32]. In the absence of coordinated management, however, severe crop damage has led to reports of unregulated persecution, including poisoning and indiscriminate killing, which can accelerate local declines without resolving conflict [5,28,29].

Our results suggest that structured removal programs anchored in continuous demographic monitoring offer a viable alternative to both inaction and uncontrolled eradication. By reducing local densities during critical periods while retaining a viable population in the landscape, such programs can simultaneously mitigate agricultural damage, generate robust population estimates, and define safe operational limits for removals, thereby supporting the long-term persistence of the species in these landscapes. In this sense, management functions not only as a tool for conflict mitigation, but also as a mechanism for learning and adaptive governance in agricultural–natural landscape mosaics.

### Population management as an effective tool for reducing crop damage

Crop damage to second-crop corn declined consistently throughout the study period and was positively associated with annual population size estimates, indicating that reductions in WLP abundance translated into lower agricultural losses (Figure 3). Although interannual variability persisted, years characterized by smaller populations generally exhibited reduced proportions of damaged corn area, supporting the premise that population management can effectively mitigate human–wildlife conflict in large-scale agricultural landscapes. However, local damage levels are shaped not only by total population size, but also by herd spatial distribution, proximity to crops during vulnerable phenological stages, and the temporal overlap between herd movements and planting or harvest schedules [30,33]. Even so, the consistent downward trend in crop damage accompanying population decline indicates that management-driven reductions in abundance alleviated pressure on crops at the regional, whole-study-area scale, overriding local stochasticity in herd behavior.

The economic relevance of this conflict is well documented. In the southern region of ENP, WLPs were estimated to damage, on average, 12.9% of corn fields annually, generating cumulative losses approaching three million US dollars for local producers [30]. Comparable economic impacts have been reported in other agricultural frontiers, including Mato Grosso state, where damage associated with WLP crop raiding also reached millions of dollars annually [29,56], underscoring the broad-scale socioeconomic implications of unmanaged populations.

Non-lethal deterrents to crop raiding animals can be an alternative to culling or removals. [56] propose non-lethal mitigation strategies, such as fencing, deterrents, and spatial avoidance, as effective tools to reduce crop raiding by forest ungulates. While these measures may be suitable for smallholdings or localized conflict scenarios [30], their applicability is limited in extensive agricultural systems where properties span thousands of hectares and damage occurs simultaneously across multiple fields. In such contexts, the logistical complexity and economic costs of field-level exclusion often exceed feasible thresholds, particularly when herds are large and highly mobile, and may also create barriers that restrict movement of other species using these landscape mosaics.

Discussions with stakeholders indicate that crop damage levels around 7% are considered economically critical for second-crop corn production (D. C. Neves, pers. comm.). After three years, our management program kept damage below the initial 12-14% levels (Table 2).

Reducing local densities in conflict areas led to measurable declines in crop damage without eliminating WLPs, balancing agricultural production and species persistence.

In addition to definitive removals, we tested an alternative strategy based on temporary removals. During two years of the program, we implemented temporary removals in which herds were captured prior to and during the planting of second-crop corn and held in captivity until the end of the critical crop period, after which individuals could be released back into the wild. This strategy was designed as an adaptive management trial to evaluate whether temporarily removing animals during peak crop vulnerability would reduce damage while maintaining population integrity. In practice, temporary removals were effective at reducing crop damage within the corresponding management year, as herds were largely absent from agricultural areas during the most vulnerable stages of crop development. However, this approach required intensive capture effort, prolonged maintenance of animals in captivity, and high operational costs, limiting its scalability relative to definitive removals. Although temporary holding also provided flexibility for future releases and adaptive decision-making, its implementation as a long-term strategy is constrained by logistical and economic considerations.

### Environmental variation as a backdrop

Our environmental analyses indicate that, within the range of variation observed in our time series, precipitation, temperature, burned area and landscape composition were not strong modulators of population dynamics or age structure when compared with the effect of management (Figure 4). Neither the annual population growth rate (rₜ) nor the proportion of juveniles showed significant relationships with environmental variables measured in year t, for rₜ, or in the previous year (t − 1), for juveniles, with low explanatory power and effect sizes generally close to zero (Table S3). Over the eight-year horizon, the main detectable demographic signal was the availability of corn as food, while removals of individuals and crop damage showed a concurrent decline and remained positively related to annual population size.

For rₜ, the clearest tendency was a decline in population growth rate with increasing annual precipitation, with a moderately negative slope, although this relationship was not statistically significant. This result may reflect sampling noise or more complex interactions among rainfall, resource availability and management logistics that are not captured by simple linear models fitted to a short series. In other ungulates, longer time series show that climate and density can interact strongly with harvest or culling to determine population growth, as in white-tailed deer (*Odocoileus virginianus*) affected simultaneously by winter severity, density and harvest intensity [16], or in moose (*Alces alces*) along bioclimatic gradients where density, habitat abundance and summer conditions explain variation in vital rates [17]. In European wild boar (*S. scrofa*), winter warming and increased food availability alter both age structure and the relative efficacy of different hunting regimes, making some types of removal more or less efficient at limiting population growth [18]. The weak climatic signal in our system suggests that, over the period analyzed, the effect of removal-based management overshadowed potential environmental effects of smaller magnitude.

In many tropical ungulate systems, rainfall is a key driver of resource availability and population dynamics, often mediating recruitment and survival through its effects on primary productivity [61]. However, in our study area, the agricultural–natural landscape mosaic likely buffers these effects by providing a relatively stable and continuous food supply across seasons and years, combining cultivated crops with native vegetation resources. This may reduce the strength of rainfall-driven fluctuations in resource availability and, consequently, dampen the demographic responses typically observed in more seasonal or resource-limited systems. Such conditions may also alter the classical population dynamics described for WLPs, in which high reproductive potential and resource-driven cycles can lead to rapid population increases followed by declines [10]. In our system, sustained high food availability combined with removal-based management may limit both strong compensatory responses and population crashes, maintaining populations at intermediate densities. However, we cannot exclude the possibility that such dynamics could emerge over longer time scales or under different environmental or management conditions.

For the proportion of juveniles, relationships with precipitation, mean temperature and burned area in t − 1 were shallow and had low R² values, indicating that interannual variation in these variables, as observed here, did not generate consistent changes in recruitment. The most structured associations involved the proportions of anthropogenic and native area in t − 1, with opposite slopes and intermediate explanatory power, although still below conventional significance thresholds. This pattern is ecologically plausible, because more agricultural land can increase food availability and potentially favors reproduction and juvenile survival, while also intensifying human disturbance and additional risks. Studies in other systems reinforce that environmental variation can modulate demography and mortality risk through shifts in habitat use and trophic interactions, such as reindeer (*Rangifer tarandus*) that are displaced into forested areas with higher predation risk under severe winter conditions [62] or cervid populations in which climate and plant productivity alter the age and sex composition of kills by predators, making mortality more or less additive among years [63].

Given the small number of years, the collinearity among land-use and land-cover metrics and our deliberately conservative choice of simple bivariate models, these results should be interpreted as exploratory indications rather than firm evidence of a lack of environmental control over WLP dynamics. In summary, in the specific context of our study, removal-based management emerges as the main driver of observed changes in abundance and population growth, whereas annual environmental fluctuations act largely as a backdrop, without producing consistent changes in rₜ or in the proportion of juveniles on their own.

### Implications for managing gregarious species in agricultural frontiers

Our results show that it’s possible to reduce the density of a social species in a tropical agricultural frontier without immediate demographic compensation through accelerated recruitment. Over eight years of management, crop damage declined, and there was a consistent positive relationship between population size and damage intensity in large-scale agricultural landscapes of the Cerrado.

For WLPs, which are simultaneously threatened and a recurrent source of conflict with agriculture in the Neotropics [29,30,51], this highlights that monitored population management can reconcile agricultural production with species persistence, maintaining viable populations while reducing conflict in highly modified landscapes. However, scaling up such interventions raises a critical and often overlooked challenge: the destination of removed individuals. In our case, more than 4,000 WLPs were absorbed by licensed commercial breeding facilities, providing a practical outlet for removed animals. Nevertheless, this solution remains incipient in Brazil and may prove challenging if population management were expanded to the scale reported in other regions, such as statewide agricultural frontiers [29,56]. Management solutions are hindered by data scarcity and the absence of wildlife management policies and regulations, particularly in Brazil, a mega-biodiverse country that is also a major worldwide producer of corn and soybeans [30,64,65,66,67]. This challenges the development of knowledge and policy solutions for coexistence.

This study provides a rare empirical foundation to inform management policies in conflict areas and advance the global debate on managing large gregarious mammals in agricultural landscapes. It documents how a real-world management program affects abundance, age structure, and crop damage in a Neotropical social ungulate using high-resolution field data.

## Acknowledgments

We are grateful to CAPES (Coordenação de Aperfeiçoamento de Pessoal de Nível Superior) for essential financial support. This research was made possible by the collaboration of the Sindicato Rural de Chapadão do Céu, Goiás, and by the engagement of rural producers operating along the southern boundary of Emas National Park. We are especially indebted to the Tropical Sustainability Institute’s field team, whose dedication in the field was fundamental for data collection under often challenging conditions. We also thank ICMBio, in particular the administration of Emas National Park, for logistical and institutional support, and CENAP for continuous technical guidance throughout the project. We are additionally grateful to Eduarda Samara for help with data entry and Leandro Scoss for his valuable support with population estimation analyses. Finally, we acknowledge producer Thomas Peixoto for his support, collaboration, and active engagement with the agricultural sector throughout this work.

## Supporting Information

### Tables

Table S1. Farm-level estimates of crop damage attributed to WLPs between 2017 and 2024. For each farm–year combination, we report the spatial resolution of the satellite imagery used (pixel size), the total damaged area (ha), the inspected corn area (ha), and the proportion of inspected area classified as damaged (%). Damage was quantified at the field level from satellite imagery acquired during the reproductive stage of second-crop corn (typically May), when intact fields exhibit high NDVI values and damaged areas are characterized by exposed soil or severely reduced vegetation cover. Within each field, damaged areas were delineated as minimum convex polygons (MCPs) encompassing contiguous patches of impacted vegetation. This MCP, and not the field, is the inspected area. Values represent the sum of inspected and damaged areas for all affected fields within each farm and year. Years marked as NA indicate periods when satellite imagery was unavailable, when second-crop corn was not planted, when fields were effectively protected from WLP access (e.g., fencing or trenches), or when no WLP damage was detected.

Table S2. Summary of WLP capture events conducted between 2017 and 2024 in the southern Cerrado, Brazil. For each capture event, the table reports the year, capture date, site (farm or protected area), specific trapping location, and number of adults (A) and juveniles (J) captured.

Table S3. Summary of univariate statistical models assessing the effects of management intensity and environmental covariates on population abundance, age structure and crop damage in the WLP population monitored. For each response–predictor combination, the table reports the intercept and slope (estimate ± standard error), degrees of freedom (df), F statistic, coefficient of determination (R²), P-value and sample size (n).

The first two models describe the relationships between population size estimated using Leslie removal model and cumulative definitive removals, and between the proportion of juveniles and cumulative removals. The third and fourth models evaluate crop damage in corn fields, relating the proportion of damaged area to (i) year (centered at 2017) and (ii) estimated population size. These models were fitted as linear mixed-effects models with farm identity included as a random effect; therefore, both marginal (R²m, variance explained by fixed effects) and conditional (R²c, variance explained by fixed and random effects combined) coefficients of determination are reported.

Subsequent models assess associations between the annual population growth rate 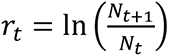 and environmental variables measured in year *t* (precipitation, mean temperature, percentage of burned area, anthropogenic cover and native cover). The final set of models relates the proportion of juveniles in year *t* to the same environmental variables measured in the previous year (*t − 1*), representing the climatic, fire and land-cover context under which the juvenile cohort was produced.

## Figures

Figure S1. Home ranges of 12 WLP herds monitored with GPS collars between 2019 and 2022 in the southern portion of ENP and surrounding agricultural lands in the Brazilian Cerrado, spanning the states of Goiás and Mato Grosso do Sul, Brazil (from [30,33]).

Polygons represent 95% AKDE home ranges estimated from GPS telemetry data [30,33] and are overlaid on satellite imagery to illustrate the spatial distribution of herds within a heterogeneous agricultural–natural mosaic.

Home ranges varied markedly in both size and landscape composition. Larger ranges were predominantly associated with native vegetation within the park, whereas smaller ranges occurred in highly anthropogenic agricultural areas. Herds occupying mixed landscapes, composed of agricultural fields interspersed with native vegetation, exhibited intermediate spatial extents.

Only herds occurring in the southern portion of the park and surrounding agricultural areas were included in the population control program.

Figure S2. Hierarchical organization of sampling units and crop damage assessment in second-crop corn in the study area. Satellite imagery (Google Earth; 26 May 2023) illustrating the spatial structure of agricultural areas used for damage assessment, based on an example. (a) Representative farm boundary (light blue polygon) encompassing multiple second-season corn fields, with field areas indicated (ha). (b) Individual corn field showing the spatial extent of damage caused by WLP, delineated as a minimum convex polygon (MCP), which defines the inspected area used for damage quantification. (c) Detailed view of damage patterns within the MCP, highlighting areas of vegetation loss associated with WLP activity.

Crop damage was calculated as the proportion of the inspected area classified as damaged, where the inspected area corresponds to the MCP. Thus, the denominator is the MCP area, and the numerator is the area of pixels classified as damaged within this polygon. Values shown in panels (b) and (c) illustrate this relationship.

Figure S3. Spatial distribution of WLP capture locations and annual study areas associated with the experimental management program conducted between 2017 and 2024 in the region surrounding ENP, central Brazilian Cerrado. Black points indicate capture locations, and grey polygons represent the annual study area defined by a 5.24 km radius buffer around each capture location. The dashed line denotes the boundary of ENP.

Figure S4. Temporal variation in demographic, environmental, disturbance, and landscape variables in the study area between 2017 and 2024. Panels show (a) estimated population size, (b) proportion of juveniles, (c) annual precipitation (mm), (d) mean annual temperature (°C), (e) proportion of area affected by fire (%), (f) crop damage in second-crop corn (%), (g) anthropogenic cover (%), and (h) native vegetation cover (%). Grey bars represent observed annual values. Blue lines represent smoothed temporal trends fitted using locally weighted regression (LOESS) to highlight general temporal patterns. Trends are shown for descriptive purposes only and should not be interpreted as formal statistical models.

