## Supplementary figures and images for "Ungulate conservation: Lessons from experimental white-lipped peccary management in agricultural-natural landscape mosaics of the Brazilian Cerrado"

### Figure S1

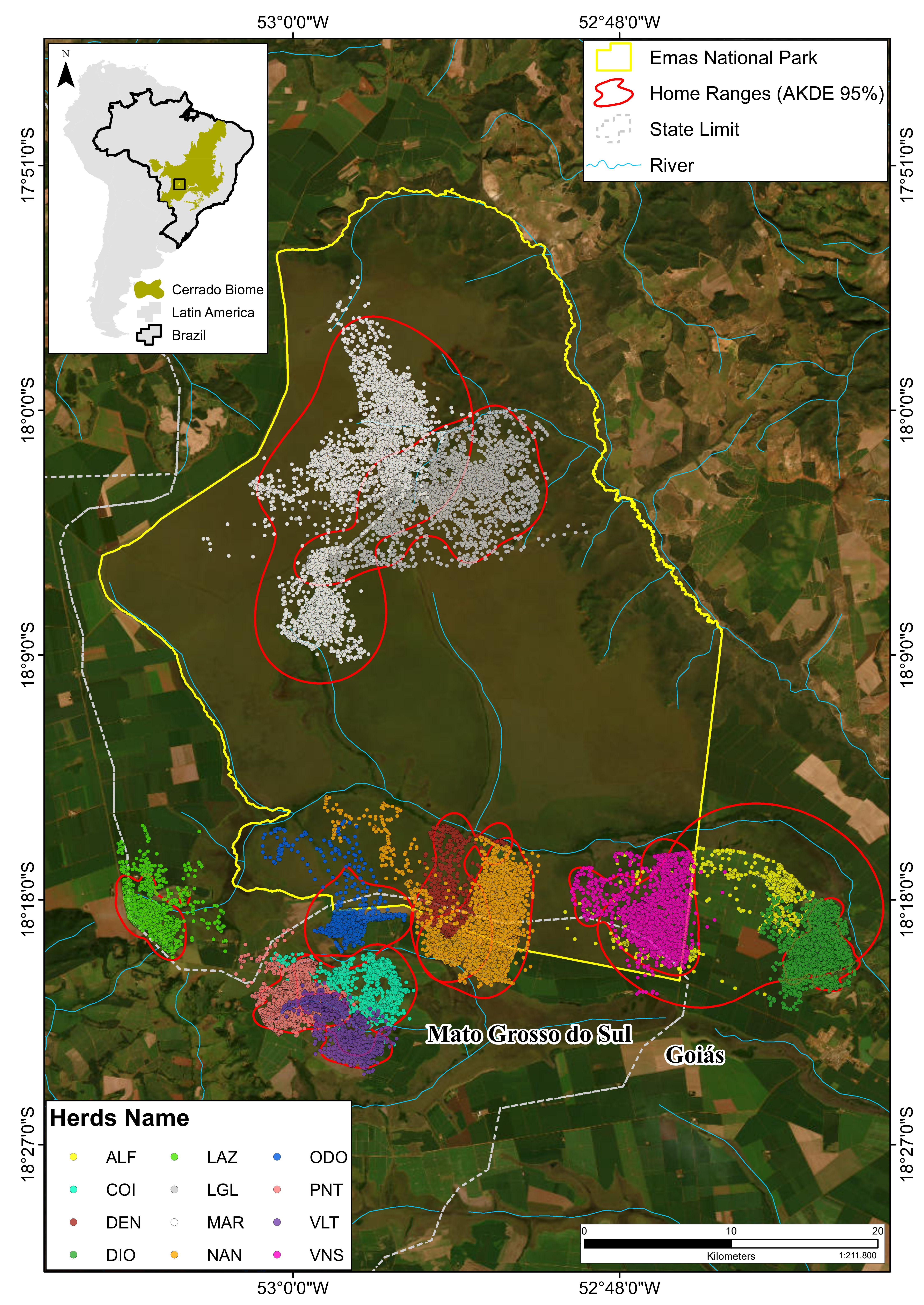

### Figure S2

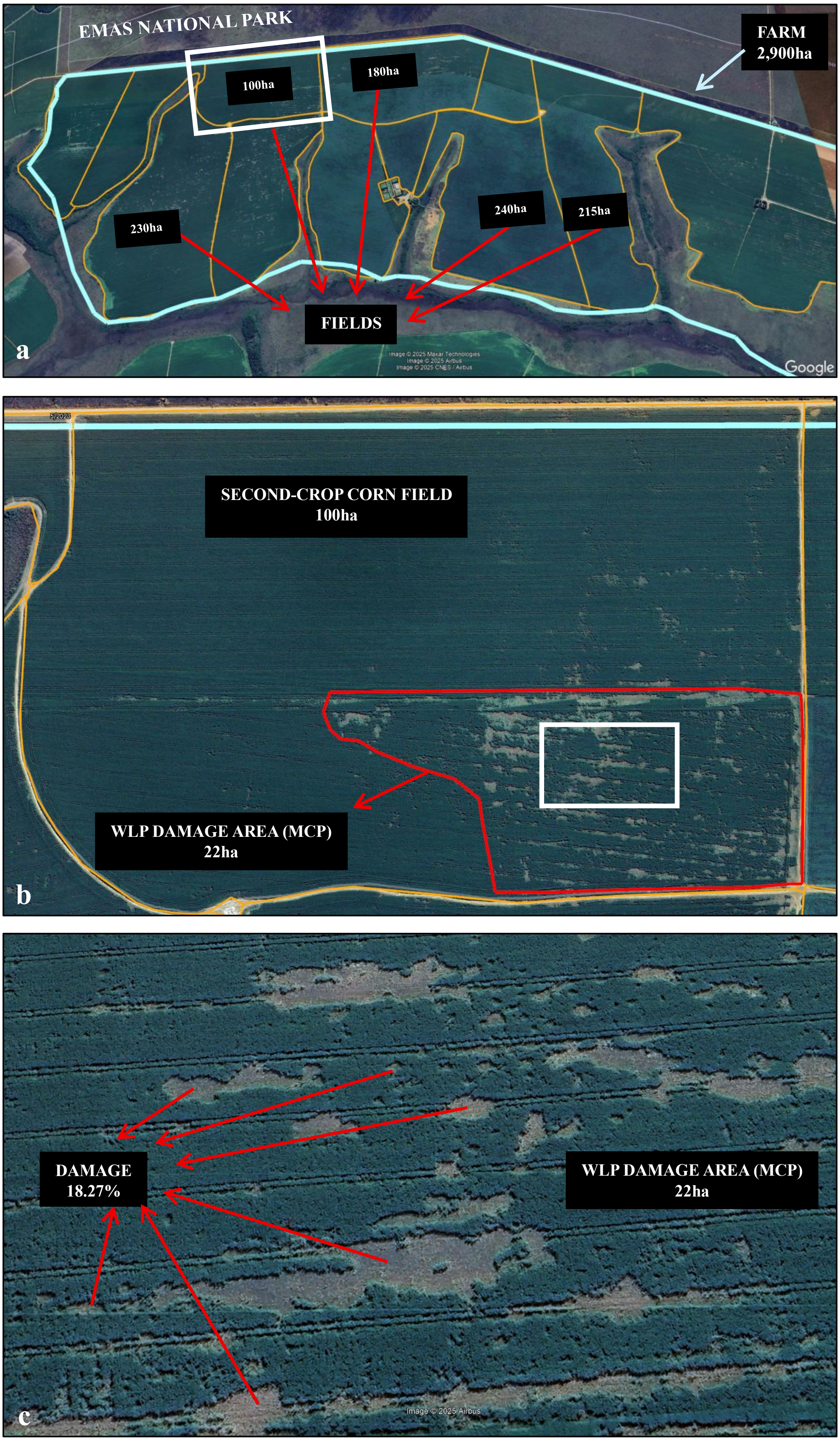

### Figure S3

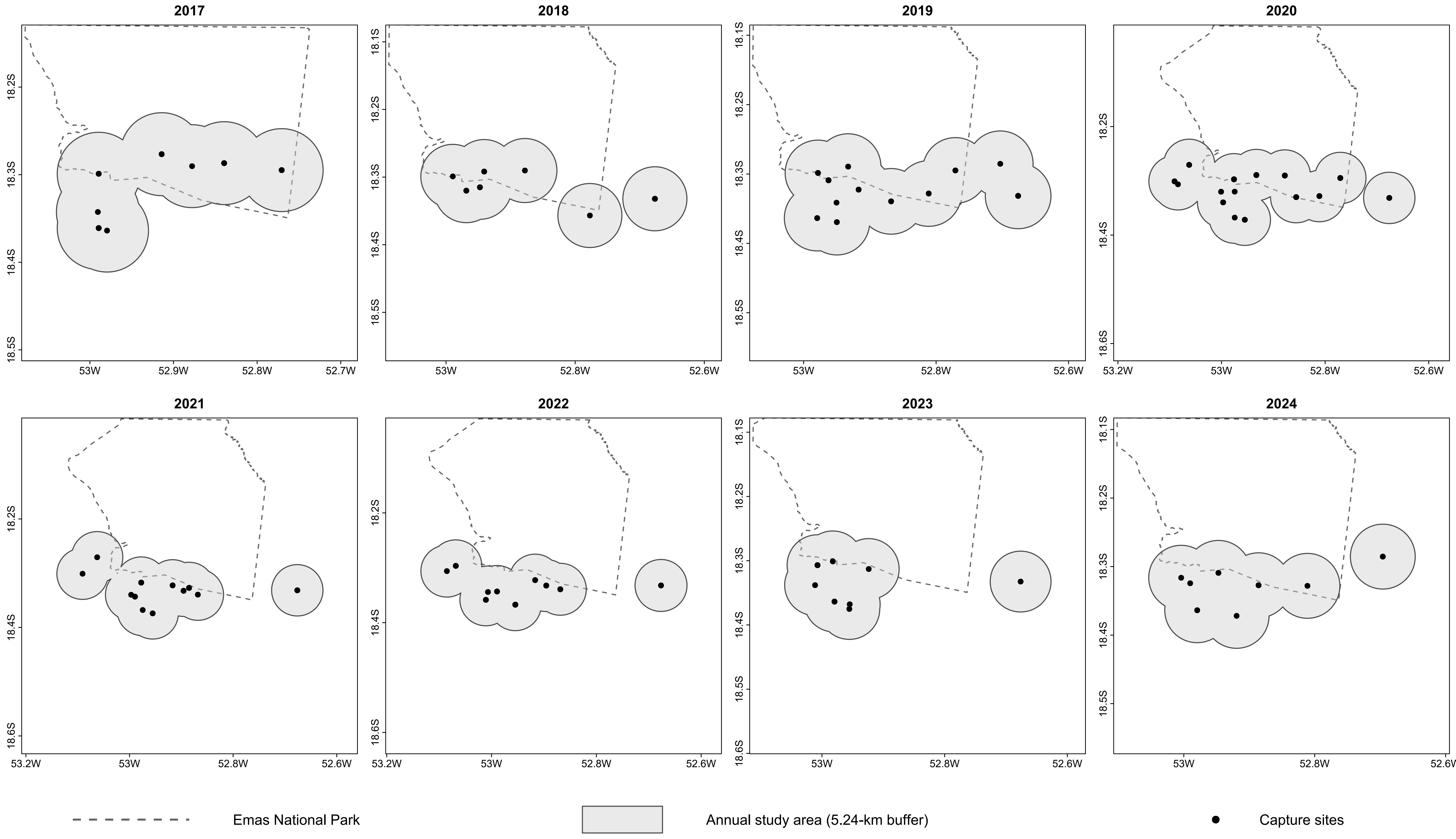

### Figure S4

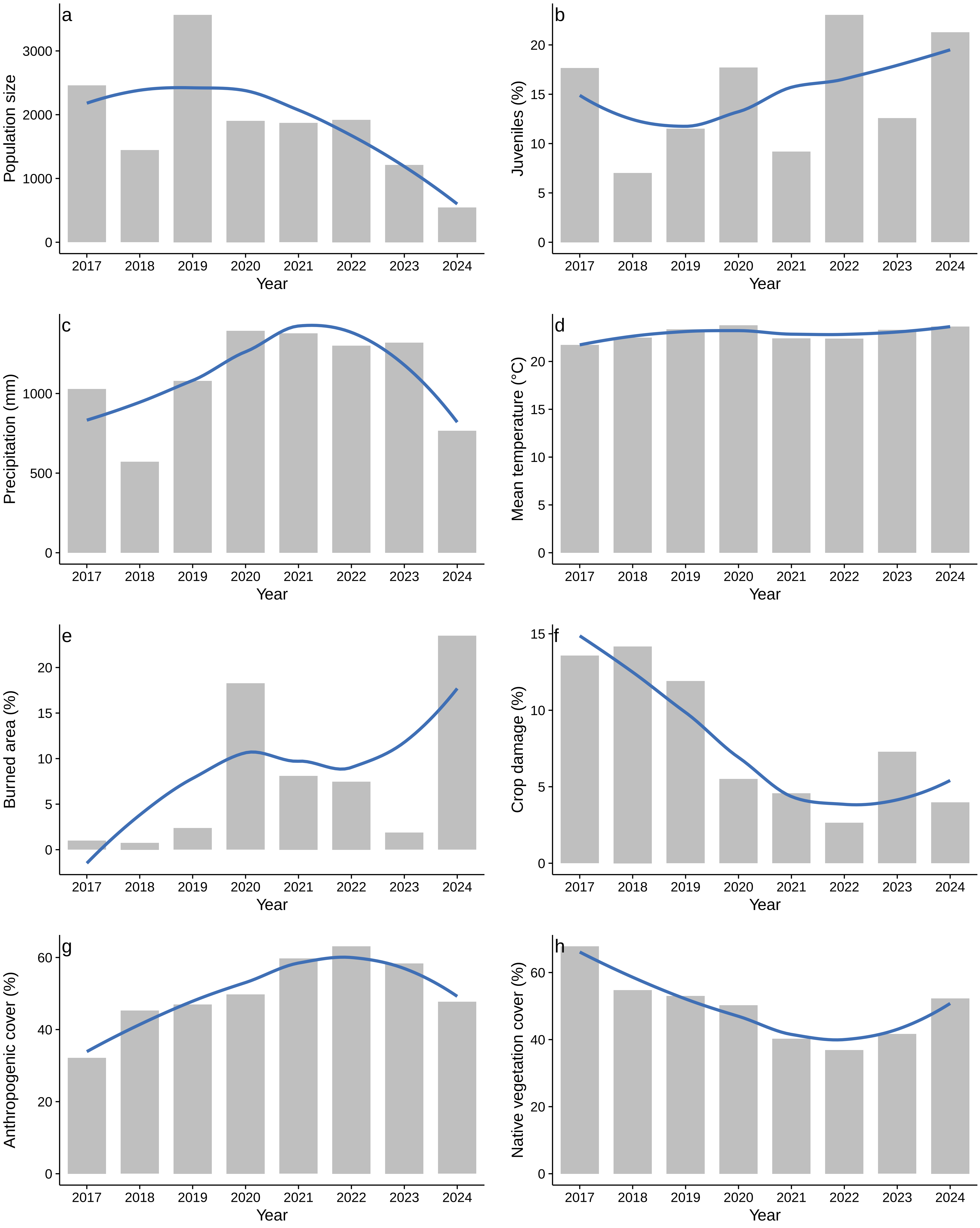
