## Supplementary material for "Ungulate conservation: Lessons from experimental white-lipped peccary management in agricultural-natural landscape mosaics of the Brazilian Cerrado": Table S1

| Farm | Year | Pixel size (m) | Damaged area (ha) | Inspected area (ha) | Damage proportion (%) |
| --- | --- | --- | --- | --- | --- |
| 01 | 2017 | 5 | 108.50 | 509.00 | 21.32 |
| 01 | 2018 | 5 | 41.60 | 360.00 | 11.56 |
| 01 | 2019 | 5 | 54.10 | 391.00 | 13.84 |
| 01 | 2020 | 2 | 13.80 | 181.00 | 7.62 |
| 01 | 2021 | 2 | 3.20 | 100.00 | 3.20 |
| 01 | 2022 | 2 | NA | NA | NA |
| 01 | 2023 | 2 | NA | NA | NA |
| 01 | 2024 | 2 | 1.00 | 39.00 | 2.56 |
| 02 | 2017 | 10 | 173.30 | 973.00 | 17.81 |
| 02 | 2018 | 10 | 376.30 | 2,060.00 | 18.27 |
| 02 | 2019 | 10 | 259.20 | 1,688.00 | 15.36 |
| 02 | 2020 | 2 | 144.10 | 1,667.00 | 8.64 |
| 02 | 2021 | 2 | 48.50 | 1,426.00 | 3.40 |
| 02 | 2022 | 2 | 29.10 | 1,149.00 | 2.53 |
| 02 | 2023 | 2 | 120.30 | 1,157.00 | 10.40 |
| 02 | 2024 | 2 | 43.60 | 893.00 | 4.88 |
| 03 | 2017 | 5 | 15.90 | 349.00 | 4.56 |
| 03 | 2018 | 5 | 17.00 | 450.00 | 3.78 |
| 03 | 2019 | 5 | NA | NA | NA |
| 03 | 2020 | 2 | 3.00 | 246.00 | 1.22 |
| 03 | 2021 | 2 | 3.00 | 129.00 | 2.33 |
| 03 | 2022 | 2 | NA | NA | NA |
| 03 | 2023 | 2 | 14.20 | 857.00 | 1.66 |
| 03 | 2024 | 2 | 7.30 | 291.00 | 2.51 |
| 04 | 2017 | 5 | 10.10 | 69.00 | 14.64 |
| 04 | 2018 | 5 | 20.70 | 136.00 | 15.22 |
| 04 | 2019 | 5 | NA | NA | NA |
| 04 | 2020 | 2 | NA | NA | NA |
| 04 | 2021 | 2 | 2.80 | 58.00 | 4.83 |
| 04 | 2022 | 2 | NA | NA | NA |
| 04 | 2023 | 2 | 10.40 | 76.00 | 13.68 |
| 04 | 2024 | 2 | 3.80 | 91.00 | 4.18 |
| 05 | 2017 | 10 | 28.40 | 445.00 | 6.38 |
| 05 | 2018 | 10 | NA | NA | NA |
| 05 | 2019 | 10 | 21.40 | 325.00 | 6.58 |
| 05 | 2020 | 2 | 28.30 | 715.00 | 3.96 |
| 05 | 2021 | 2 | 57.30 | 848.00 | 6.76 |
| 05 | 2022 | 2 | 23.60 | 695.00 | 3.40 |
| 05 | 2023 | 2 | 33.60 | 414.00 | 8.12 |
| 05 | 2024 | 2 | 15.70 | 188.00 | 8.35 |
| 06 | 2017 | 10 | 24.00 | 305.00 | 7.87 |
| 06 | 2018 | 10 | 8.40 | 315.00 | 2.67 |
| 06 | 2019 | 10 | 11.30 | 548.00 | 2.06 |
| 06 | 2020 | 2 | 27.40 | 1,133.00 | 2.42 |
| 06 | 2021 | 2 | 32.70 | 932.00 | 3.51 |
| 06 | 2022 | 2 | 3.80 | 445.00 | 0.85 |
| 06 | 2023 | 2 | 2.20 | 27.00 | 8.15 |
| 06 | 2024 | 2 | 3.50 | 394.00 | 0.89 |
| 07 | 2017 | 10 | NA | NA | NA |
| 07 | 2018 | 10 | NA | NA | NA |
| 07 | 2019 | 10 | 7.60 | 20.00 | 38.00 |
| 07 | 2020 | 2 | NA | NA | NA |
| 07 | 2021 | 2 | 24.50 | 188.00 | 13.03 |
| 07 | 2022 | 2 | 6.50 | 147.00 | 4.42 |
| 07 | 2023 | 2 | 11.20 | 113.00 | 9.91 |
| 07 | 2024 | 2 | 3.00 | 60.00 | 5.00 |
| 08 | 2017 | 5 | 0.60 | 29.00 | 2.07 |
| 08 | 2018 | 5 | NA | NA | NA |
| 08 | 2019 | 5 | NA | NA | NA |
| 08 | 2020 | 2 | 1.00 | 7.00 | 14.29 |
| 08 | 2021 | 2 | 3.20 | 157.00 | 2.04 |
| 08 | 2022 | 2 | 1.50 | 7.00 | 21.43 |
| 08 | 2023 | 2 | 3.30 | 35.00 | 9.43 |
| 08 | 2024 | 2 | NA | NA | NA |
