## Supplementary material for "Ungulate conservation: Lessons from experimental white-lipped peccary management in agricultural-natural landscape mosaics of the Brazilian Cerrado": Table S2

| Year | Capture date | Site | Trap site | A | J |
| --- | --- | --- | --- | --- | --- |
| 2017 | 2017-07-11 | Farm 07 | Base Gloria | 26 | 1 |
| 2017 | 2017-08-06 | Farm 05 | Brete 1 COI | 44 | 10 |
| 2017 | 2017-08-08 | Emas National Park | VII | 65 | 2 |
| 2017 | 2017-08-20 | Farm 05 | Brete 2 | 61 | 7 |
| 2017 | 2017-09-07 | Farm 07 | Base Gloria | 23 | 9 |
| 2017 | 2017-09-07 | Emas National Park | VII | 28 | 5 |
| 2017 | 2017-09-08 | Farm 07 | Base Gloria | 26 | 8 |
| 2017 | 2017-09-12 | Emas National Park | III | 22 | 0 |
| 2017 | 2017-09-13 | Farm 07 | Base Gloria | 12 | 2 |
| 2017 | 2017-09-28 | Farm 07 | Base Gloria | 58 | 2 |
| 2017 | 2017-10-07 | Farm 07 | Base Gloria | 41 | 0 |
| 2017 | 2017-10-09 | Emas National Park | III | 46 | 4 |
| 2017 | 2017-10-10 | Farm 05 | Brete 03 | 37 | 15 |
| 2017 | 2017-10-13 | Emas National Park | I | 23 | 11 |
| 2017 | 2017-10-14 | Emas National Park | III | 17 | 6 |
| 2017 | 2017-10-26 | Farm 05 | Brete 03 | 52 | 17 |
| 2017 | 2017-10-28 | Emas National Park | III | 41 | 14 |
| 2017 | 2017-11-01 | Farm 05 | Brete 03 | 28 | 17 |
| 2017 | 2017-11-04 | Emas National Park | VII | 40 | 14 |
| 2017 | 2017-11-06 | Emas National Park | II | 19 | 6 |
| 2017 | 2017-11-07 | Emas National Park | VII | 37 | 6 |
| 2017 | 2017-11-15 | Emas National Park | II | 46 | 19 |
| 2017 | 2017-11-29 | Farm 05 | Brete 03 | 9 | 3 |
| 2017 | 2017-11-29 | Emas National Park | VII | 20 | 1 |
| 2017 | 2017-12-02 | Emas National Park | II | 18 | 4 |
| 2017 | 2017-12-13 | Farm 05 | Brete 03 | 19 | 1 |
| 2018 | 2018-04-21 | Emas National Park | Brete do Gloria | 16 | 2 |
| 2018 | 2018-04-24 | Farm 07 | Base Gloria | 46 | 11 |
| 2018 | 2018-04-27 | Emas National Park | Brete do Gloria | 39 | 7 |
| 2018 | 2018-05-02 | Emas National Park | Brete do Gloria | 44 | 0 |
| 2018 | 2018-05-11 | Emas National Park | I | 70 | 5 |
| 2018 | 2018-05-21 | Farm 08 | Brete Matinha | 74 | 3 |
| 2018 | 2018-06-23 | Farm 03 | Ceva Cascalheira | 25 | 0 |
| 2018 | 2018-06-27 | Farm 08 | Brete Matinha | 22 | 8 |
| 2018 | 2018-07-03 | Farm 08 | Brete Matinha | 20 | 0 |
| 2018 | 2018-07-06 | Farm 08 | Brete Matinha | 18 | 0 |
| 2018 | 2018-07-14 | Farm 02 | Brete 01 | 18 | 0 |
| 2018 | 2018-07-18 | Farm 02 | Brete 01 | 36 | 3 |
| 2018 | 2018-07-26 | Farm 02 | Brete 01 | 54 | 3 |
| 2018 | 2018-08-08 | Farm 02 | Brete 02 | 69 | 12 |
| 2018 | 2018-08-16 | Farm 02 | Brete 02 | 86 | 0 |
| 2018 | 2018-09-05 | Farm 02 | Brete 02 | 79 | 0 |
| 2019 | 2018-11-28 | Farm 08 | Brete Matinha | 25 | 9 |
| 2019 | 2018-12-04 | Farm 08 | Brete Matinha | 27 | 5 |
| 2019 | 2018-12-05 | Emas National Park | Ceva da Onça | 20 | 10 |
| 2019 | 2018-12-07 | Emas National Park | Ceva da Onça | 29 | 9 |
| 2019 | 2018-12-12 | Farm 08 | Brete Matinha | 16 | 0 |
| 2019 | 2018-12-19 | Emas National Park | VII | 45 | 12 |
| 2019 | 2019-01-08 | Emas National Park | VII | 60 | 12 |
| 2019 | 2019-01-31 | Farm 08 | Brete Matinha | 32 | 4 |
| 2019 | 2019-02-03 | Farm 07 | Bosque | 40 | 3 |
| 2019 | 2019-02-04 | Farm 07 | Bosque | 30 | 3 |
| 2019 | 2019-02-05 | Farm 05 | I | 26 | 2 |
| 2019 | 2019-02-06 | Farm 07 | Bosque | 60 | 2 |
| 2019 | 2019-02-15 | Farm 05 | Cana sede | 32 | 5 |
| 2019 | 2019-02-16 | Farm 05 | Cana sede | 39 | 8 |
| 2019 | 2019-03-12 | Farm 08 | Cana graxa | 46 | 11 |
| 2019 | 2019-03-13 | Farm 05 | Vereda / Dreno | 42 | 5 |
| 2019 | 2019-03-18 | Emas National Park | Glória 2 | 16 | 0 |
| 2019 | 2019-03-19 | Emas National Park | Glória 2 | 66 | 8 |
| 2019 | 2019-04-26 | Farm 02 | Ceva do Portão | 32 | 0 |
| 2019 | 2019-05-03 | Farm 05 | I | 27 | 5 |
| 2019 | 2019-05-04 | Farm 02 | Ceva do Portão | 22 | 3 |
| 2019 | 2019-05-08 | Farm 01 | Ceva do Brejo | 21 | 0 |
| 2019 | 2019-05-09 | Farm 02 | Ceva do Brejo | 12 | 0 |
| 2019 | 2019-05-13 | Farm 02 | Ceva do Brejo | 20 | 4 |
| 2019 | 2019-05-14 | Farm 01 | Ceva do Brejo | 25 | 0 |
| 2019 | 2019-05-14 | Farm 02 | Ceva do Brejo | 6 | 0 |
| 2019 | 2019-05-15 | Farm 02 | Ceva do Brejo | 6 | 0 |
| 2019 | 2019-05-18 | Farm 02 | Ceva do Brejo | 21 | 0 |
| 2019 | 2019-05-20 | Farm 02 | Ceva do Brejo | 19 | 0 |
| 2019 | 2019-05-22 | Farm 02 | Ceva do Brejo | 16 | 2 |
| 2019 | 2019-05-24 | Farm 02 | Ceva do Portão | 17 | 1 |
| 2019 | 2019-05-30 | Farm 02 | Ceva do Brejo | 36 | 0 |
| 2019 | 2019-06-05 | Farm 02 | Ceva do Portão | 35 | 4 |
| 2019 | 2019-06-11 | Farm 02 | Ceva do Brejo | 27 | 2 |
| 2020 | 2019-12-30 | Emas National Park | ARU | 14 | 4 |
| 2020 | 2019-12-31 | Emas National Park | ARU | 17 | 0 |
| 2020 | 2019-12-31 | Emas National Park | Ceva da Onça | 18 | 2 |
| 2020 | 2020-01-03 | Emas National Park | Ceva da Onça | 23 | 11 |
| 2020 | 2020-01-07 | Emas National Park | ARU | 8 | 4 |
| 2020 | 2020-01-09 | Emas National Park | I | 17 | 2 |
| 2020 | 2020-01-11 | Emas National Park | I | 15 | 2 |
| 2020 | 2020-01-11 | Emas National Park | VII | 14 | 0 |
| 2020 | 2020-01-14 | Emas National Park | VII | 10 | 3 |
| 2020 | 2020-01-16 | Emas National Park | Glória 3 | 25 | 0 |
| 2020 | 2020-01-18 | Emas National Park | Glória 2 | 21 | 2 |
| 2020 | 2020-01-18 | Emas National Park | Glória 3 | 10 | 6 |
| 2020 | 2020-01-19 | Emas National Park | VII | 23 | 0 |
| 2020 | 2020-01-20 | Emas National Park | Glória 2 | 13 | 1 |
| 2020 | 2020-01-20 | Emas National Park | Glória 3 | 24 | 3 |
| 2020 | 2020-01-20 | Emas National Park | I | 16 | 7 |
| 2020 | 2020-01-21 | Emas National Park | Glória 2 | 11 | 2 |
| 2020 | 2020-01-21 | Emas National Park | VII | 28 | 12 |
| 2020 | 2020-01-23 | Emas National Park | VII | 20 | 1 |
| 2020 | 2020-01-25 | Emas National Park | VII | 22 | 8 |
| 2020 | 2020-01-26 | Emas National Park | Glória 2 | 19 | 4 |
| 2020 | 2020-01-27 | Emas National Park | Glória 2 | 16 | 1 |
| 2020 | 2020-01-27 | Emas National Park | I | 31 | 3 |
| 2020 | 2020-01-28 | Emas National Park | I | 31 | 13 |
| 2020 | 2020-01-29 | Farm 06 | Cana | 7 | 1 |
| 2020 | 2020-01-31 | Farm 08 | Brete Matinha | 20 | 10 |
| 2020 | 2020-01-31 | Emas National Park | I | 21 | 9 |
| 2020 | 2020-02-01 | Farm 08 | Brete Matinha | 8 | 0 |
| 2020 | 2020-02-02 | Farm 06 | Mata | 19 | 0 |
| 2020 | 2020-02-04 | Emas National Park | VII | 15 | 10 |
| 2020 | 2020-02-05 | Farm 06 | Mata | 5 | 0 |
| 2020 | 2020-02-06 | Farm 08 | Brete Matinha | 29 | 12 |
| 2020 | 2020-02-07 | Farm 05 | 3 divisas | 21 | 2 |
| 2020 | 2020-02-08 | Emas National Park | Glória 2 | 14 | 3 |
| 2020 | 2020-02-09 | Emas National Park | Glória 2 | 14 | 3 |
| 2020 | 2020-02-12 | Farm 05 | 3 divisas | 16 | 2 |
| 2020 | 2020-02-12 | Emas National Park | ARU | 37 | 4 |
| 2020 | 2020-02-13 | Farm 06 | Mata | 28 | 4 |
| 2020 | 2020-02-14 | Emas National Park | ARU | 16 | 2 |
| 2020 | 2020-02-15 | Emas National Park | Glória 3 | 19 | 1 |
| 2020 | 2020-02-16 | Farm 05 | Cana | 29 | 4 |
| 2020 | 2020-02-18 | Emas National Park | Glória 3 | 20 | 0 |
| 2020 | 2020-02-19 | Farm 07 | Cana Glória Vala | 17 | 5 |
| 2020 | 2020-02-19 | Emas National Park | Glória 3 | 12 | 3 |
| 2020 | 2020-02-20 | Farm 07 | Cana Glória Vala | 17 | 3 |
| 2020 | 2020-02-20 | Farm 06 | Cana | 8 | 0 |
| 2020 | 2020-02-25 | Farm 05 | Cana | 17 | 1 |
| 2020 | 2020-02-26 | Farm 05 | Cana | 45 | 13 |
| 2020 | 2020-02-27 | Farm 05 | Cana | 31 | 4 |
| 2020 | 2020-03-03 | Farm 05 | 3 divisas | 9 | 6 |
| 2020 | 2020-03-03 | Emas National Park | Glória 3 | 8 | 0 |
| 2020 | 2020-05-07 | Farm 05 | Gambi | 36 | 13 |
| 2020 | 2020-05-08 | Farm 06 | Lázaro | 27 | 2 |
| 2020 | 2020-05-15 | Farm 02 | II | 30 | 16 |
| 2021 | 2021-02-06 | Farm 05 | Gambi | 38 | 4 |
| 2021 | 2021-02-07 | Farm 08 | Brete Matinha | 12 | 0 |
| 2021 | 2021-02-19 | Farm 08 | Brete Matinha | 37 | 6 |
| 2021 | 2021-02-24 | Farm 01 | Ceva do Brejo | 17 | 0 |
| 2021 | 2021-02-28 | Farm 02 | Lagherta | 14 | 2 |
| 2021 | 2021-03-04 | Farm 05 | 3 divisas | 9 | 2 |
| 2021 | 2021-03-05 | Farm 06 | Mata | 20 | 5 |
| 2021 | 2021-03-06 | Farm 02 | Lagherta | 9 | 3 |
| 2021 | 2021-03-13 | Farm 02 | Lagherta | 12 | 0 |
| 2021 | 2021-03-13 | Farm 05 | Cana/soja | 7 | 0 |
| 2021 | 2021-03-15 | Farm 05 | Cana/soja | 9 | 0 |
| 2021 | 2021-03-16 | Farm 02 | Lagherta | 18 | 4 |
| 2021 | 2021-03-23 | Farm 05 | COI | 14 | 0 |
| 2021 | 2021-03-29 | Farm 02 | Curva | 14 | 0 |
| 2021 | 2021-03-30 | Farm 02 | Ceva do Portão | 33 | 4 |
| 2021 | 2021-03-31 | Farm 02 | Curva | 14 | 0 |
| 2021 | 2021-04-13 | Farm 05 | COI | 28 | 6 |
| 2021 | 2021-04-13 | Farm 08 | Brete Matinha | 16 | 0 |
| 2021 | 2021-04-16 | Farm 02 | Curva | 41 | 1 |
| 2021 | 2021-04-21 | Farm 05 | COI | 30 | 3 |
| 2021 | 2021-04-27 | Farm 02 | Ceva do Portão | 14 | 1 |
| 2021 | 2021-05-05 | Farm 02 | Ceva do PK | 12 | 1 |
| 2021 | 2021-05-06 | Farm 02 | Ceva do PK | 19 | 0 |
| 2021 | 2021-05-10 | Farm 02 | Ceva do PK | 19 | 2 |
| 2021 | 2021-05-19 | Farm 06 | Cana | 21 | 3 |
| 2021 | 2021-06-04 | Farm 02 | Curva | 17 | 3 |
| 2022 | 2022-01-25 | Farm 05 | Sede/cana/soja | 10 | 0 |
| 2022 | 2022-01-26 | Farm 02 | Curva | 16 | 2 |
| 2022 | 2022-01-27 | Farm 08 | Brete Matinha | 35 | 7 |
| 2022 | 2022-01-28 | Farm 05 | Sede/cana/soja | 22 | 4 |
| 2022 | 2022-02-05 | Farm 06 | Triângulo | 14 | 1 |
| 2022 | 2022-02-09 | Farm 05 | COI | 11 | 1 |
| 2022 | 2022-02-10 | Farm 06 | Triângulo | 23 | 2 |
| 2022 | 2022-02-12 | Farm 07 | Cana | 21 | 2 |
| 2022 | 2022-02-19 | Farm 05 | Sede/cana/soja | 59 | 16 |
| 2022 | 2022-02-22 | Farm 05 | Sede/cana/soja | 10 | 7 |
| 2022 | 2022-02-28 | Farm 05 | Sede/cana/soja | 9 | 1 |
| 2022 | 2022-03-01 | Farm 07 | 3B | 33 | 15 |
| 2022 | 2022-03-03 | Farm 06 | Mata | 29 | 10 |
| 2022 | 2022-03-05 | Farm 05 | Sede/cana/soja | 34 | 17 |
| 2022 | 2022-03-10 | Farm 08 | Brete Matinha | 10 | 1 |
| 2022 | 2022-03-14 | Farm 02 | Curva | 40 | 10 |
| 2022 | 2022-03-28 | Farm 07 | Cana | 31 | 21 |
| 2022 | 2022-03-29 | Farm 06 | Triângulo | 31 | 10 |
| 2022 | 2022-04-15 | Farm 02 | Curva | 25 | 5 |
| 2022 | 2022-04-16 | Farm 02 | Ceva do Portão | 36 | 9 |
| 2022 | 2022-04-22 | Farm 02 | Curva | 17 | 3 |
| 2022 | 2022-05-04 | Farm 02 | Ceva do Portão | 12 | 10 |
| 2022 | 2022-05-11 | Farm 01 | Murici | 17 | 1 |
| 2022 | 2022-05-13 | Farm 01 | Murici | 22 | 0 |
| 2022 | 2022-05-16 | Farm 07 | Cana | 24 | 7 |
| 2022 | 2022-05-17 | Farm 05 | COI | 10 | 6 |
| 2022 | 2022-05-19 | Farm 07 | Cana | 17 | 12 |
| 2022 | 2022-05-19 | Farm 01 | Murici | 18 | 4 |
| 2022 | 2022-05-24 | Farm 01 | Murici | 11 | 3 |
| 2022 | 2022-05-24 | Farm 02 | Ceva do Portão | 14 | 5 |
| 2022 | 2022-05-31 | Farm 01 | Murici | 21 | 5 |
| 2022 | 2022-06-02 | Farm 05 | COI | 26 | 9 |
| 2022 | 2022-06-14 | Farm 05 | Sede/cana/soja | 13 | 10 |
| 2023 | 2023-02-08 | Farm 07 | Brete da vala | 16 | 0 |
| 2023 | 2023-02-09 | Farm 07 | Brete da vala | 44 | 5 |
| 2023 | 2023-02-12 | Farm 05 | Brete do Gambi | 43 | 7 |
| 2023 | 2023-02-20 | Farm 07 | Brete da vala | 40 | 2 |
| 2023 | 2023-03-07 | Farm 07 | Brete da vala | 12 | 3 |
| 2023 | 2023-03-09 | Farm 05 | Brete cana/sede | 20 | 9 |
| 2023 | 2023-03-13 | Farm 05 | Brete do Gambi | 45 | 17 |
| 2023 | 2023-03-25 | Farm 05 | Brete do Gambi | 22 | 5 |
| 2023 | 2023-03-28 | Farm 07 | Brete da vala | 34 | 2 |
| 2023 | 2023-04-14 | Farm 05 | Brete do Gambi | 28 | 8 |
| 2023 | 2023-04-20 | Farm 07 | Brete do terreiro | 21 | 0 |
| 2023 | 2023-04-21 | Farm 07 | Brete do terreiro | 34 | 0 |
| 2023 | 2023-04-29 | Farm 05 | Brete da cana/cascalheira | 43 | 12 |
| 2023 | 2023-05-01 | Farm 08 | Brete Matinha | 21 | 0 |
| 2023 | 2023-05-08 | Farm 07 | Brete da Vagem | 27 | 0 |
| 2023 | 2023-05-08 | Farm 05 | Brete do Gambi | 31 | 0 |
| 2023 | 2023-05-13 | Farm 05 | Brete do Gambi | 25 | 4 |
| 2023 | 2023-05-18 | Farm 02 | Brete do Calcário | 15 | 1 |
| 2024 | 2024-04-03 | Emas National Park | Ceva da Onça | 24 | 13 |
| 2024 | 2024-04-09 | Farm 02 | Ceva do PK | 39 | 11 |
| 2024 | 2024-04-11 | Farm 05 | Ceva do 22 | 22 | 6 |
| 2024 | 2024-04-18 | Farm 02 | Ceva do PK | 21 | 4 |
| 2024 | 2024-04-20 | Farm 02 | Ceva do algodão | 27 | 6 |
| 2024 | 2024-04-23 | Farm 02 | Ceva do algodão | 21 | 4 |
| 2024 | 2024-04-25 | Farm 02 | Ceva do PK | 17 | 0 |
| 2024 | 2024-05-07 | Farm 07 | Ceva M3 | 14 | 6 |
| 2024 | 2024-06-21 | Farm 07 | Ceva do adubo | 13 | 0 |
| 2024 | 2024-07-09 | Farm 05 | Ceva do gambi | 39 | 19 |
| 2024 | 2024-07-10 | Farm 05 | Ceva do gambi | 8 | 0 |
| 2024 | 2024-07-11 | Farm 08 | Ceva do brejo | 9 | 0 |
| 2024 | 2024-07-12 | Farm 08 | Ceva do brejo | 16 | 4 |
