## Supplementary material for "Ungulate conservation: Lessons from experimental white-lipped peccary management in agricultural-natural landscape mosaics of the Brazilian Cerrado": Table S3

| Response | Predictor | Intercept  (± SE) | Slope  (± SE) | df | F | R² | P | n |
| --- | --- | --- | --- | --- | --- | --- | --- | --- |
| Population size (N) | Cumulative removals | 2,708.25  (± 449.23) | -0.377  (± 0.167) | 1, 6 | 5.12 | 0.46 | 0.0643 | 8 |
| Juveniles (%) | Cumulative removals | 11.80  (± 3.63) | 0.001  (± 0.001) | 1, 6 | 1.13 | 0.158 | 0.33 | 8 |
| Crop damage (%) | Year (centered at 2017) | 0.119  (± 0.019) | -0.010  (± 0.004) | 1, 41 | 7.17 | 0.111m  0.286c | 0.0106 | 50 |
| Crop damage (%) | Population size (N) | 0.082  (± 0.013) | 0.022  (± 0.009) | 1, 41 | 6.07 | 0.099m  0.242c | 0.018 | 50 |
| Growth rate (r) | Precipitation (mm) | 1.15  (± 0.84) | -0.001  (± 0.001) | 1, 5 | 2.80 | 0.359 | 0.155 | 7 |
| Growth rate (r) | Mean temperature (°C) | 2.19  (± 8.19) | -0.105  (± 0.359) | 1, 5 | 0.09 | 0.017 | 0.781 | 7 |
| Growth rate (r) | Burned area (%) | -0.27  (± 0.33) | 0.009  (± 0.041) | 1, 5 | 0.05 | 0.01 | 0.831 | 7 |
| Growth rate (r) | Anthropogenic area (%) | 0.10  (± 1.25) | -0.006  (± 0.024) | 1, 5 | 0.06 | 0.013 | 0.809 | 7 |
| Growth rate (r) | Native area (%) | -0.52  (± 1.22) | 0.006  (± 0.024) | 1, 5 | 0.06 | 0.013 | 0.809 | 7 |
| Juveniles (%) | Precipitation in year t-1 (mm) | 5.85  (± 10.33) | 0.008  (± 0.009) | 1, 5 | 0.76 | 0.132 | 0.423 | 7 |
| Juveniles (%) | Mean temperature in year t-1 (°C) | -32.57  (± 84.65) | 2.071  (± 3.714) | 1, 5 | 0.31 | 0.059 | 0.601 | 7 |
| Juveniles (%) | Burned area in year t-1 (%) | 15.47  (± 3.51) | -0.150  (± 0.430) | 1, 5 | 0.12 | 0.024 | 0.741 | 7 |
| Juveniles (%) | Anthropogenic area in year t-1 (%) | -4.25  (± 10.19) | 0.372  (± 0.197) | 1, 5 | 3.56 | 0.416 | 0.118 | 7 |
| Juveniles (%) | Native area in year t-1 (%) | 32.92  (± 9.89) | -0.372  (± 0.197) | 1, 5 | 3.56 | 0.416 | 0.118 | 7 |
